# Correlation of drug resistance with Single Nucleotide Variations through genome analysis and experimental validation in a multi-drug resistant clinical isolate of *M.tuberculosis*

**DOI:** 10.1101/2020.03.07.981878

**Authors:** Kausik Bhattacharyya, Vishal Nemaysh, Monika Joon, Ramendra Pratap, Mandira Varma-Basil, Mridula Bose, Vani Brahmachari

## Abstract

**Background:** Genome sequencing and genetic polymorphism analysis of clinical isolates of *M.tuberculosis* is carried out to gain further insight into molecular pathogenesis and host-pathogen interaction. Therefore the functional evaluation of the effect of single nucleotide variation (SNV) is essential. At the same time, the identification of invariant sequences unique to *M.tuberculosis* contributes to infection detection by sensitive methods. In the present study, genome analysis is accompanied by evaluation of the functional implication of the SNVs in a MDR clinical isolate VPCI591.

**Result:** By sequencing and comparative analysis of VPCI591 genome with 1553 global clinical isolates of *M.tuberculosis*, we identified 143 unique strain specific SNVs. A novel intergenic variation in VPCI591 in the putative promoter/regulatory region mapping between *embC (Rv3793)* and *embA* (*Rv3794*) genes was found to enhance the expression of *embAB*, which correlates with the high resistance of the VPCI591 to ethambutol. Similarly, the unique combination of three genic SNVs in RNA polymerase β gene (*rpoB*) in VPCI591 was evaluated for its effect on rifampicin resistance through molecular docking analysis. The comparative genomics also showed that along with variations, there are genes that remain invariant. 173 such genes were identified in our analysis.

**Conclusion:** We have demonstrated that SNVs are not always benign and can also have functional effect. We show that variations bring about quantitative changes in transcription. Our results show the collective effect of SNVs on the structure of protein, impacting the interaction between the target protein and the drug molecule, with *rpoB* as an example.

## Background

In spite of the worldwide efforts to combat mycobacterial diseases, it continues to be a great challenge to achieve this goal. In addition to the various strategies adopted by this pathogen to escape host immune system, *M.tuberculosis* has gainfully utilized genetic variability for its highly successful growth, pathogenesis, immunity and persistence (1). The complete genome sequence of the strain of *M. tuberculosis* H37Rv and the recent surge in data on clinical isolates permits high-throughput whole-genome analysis, relationship and correlation with drug resistance (2-9). This has revealed local, global and patient specific heterogeneity in *M.tuberculosis* strains (10). The increased genetic diversity and variation in *M.tuberculosis* is implicated in strain specific immunogenicity and pathogenicity (11). A large number and the most frequent occurrence of variation is seen in the genes for lipid metabolism and *PE*/*PPE* genes (12, 13).

The whole genome sequence of more than 11,000 clinical isolates of *M.tuberculosis* are reported in NCBI-SRA, however, the correlation between genetic alterations and the phenotype is carried out mainly in the drug resistance genes (14). Thus, drug resistance in tuberculosis is a phenomenon more complex than previously assumed where the association of intergenic regions and the identification of new genes have resulted from whole-genome sequence based approach (15).

We analysed the genetic polymorphism in an Indian, multi-drug resistant (MDR) clinical isolate VPCI591, from a 50 year old male patient from Vallabhbhai Patel Chest Institute (VPCI), Delhi. It is TBd-1 positive and EAI clade spoligotype and the isolate is resistant to all the first-line tuberculosis drugs (16, 17). The availability of this strain made it possible to validate selected variations and investigate their effect on the function of the gene. We carried out the comparative analysis of the variations detected in VPCI591 with the genome sequence of about 1553 global clinical isolates, to identify the shared and unique Single Nucleotide Variations (SNV). This analysis also indicated the invariant regions/genes that could be of potential use as markers for *M.tuberculosis* complex.

## Methods

### Bacterial strains and culture conditions

*M.tuberculosis* clinical isolate VPCI591 and *M.tuberculosis* H37Rv were grown at 37°C in Middlebrook 7H9 broth (Becton Dickinson) supplemented with 10 % OADC (Oleic acid, Bovine albumin fraction V, dextrose, catalase) from Himedia (FD329), with 0.2% glycerol (SRL) and 0.05% tween 80 (Sigma).

### Genome sequencing and pipeline for analysis of genome variation

Genomic DNA was isolated using standard protocol (18). The genome sequencing was done using Illumina GAIIx analyser at CSIR-Institute of genomics and Integrative biology (IGIB). TruSeq2_SE adapters were used for sequencing. Sequence read file for the clinical isolate VPCI591 have been deposited in SRA format, NCBI: SRX5802345, Bioproject accession number PRJNA540936, BioSample accession numbers: SAMN11568242.

FastQC (0.11.5) was used for deciphering overall quality statistics of the data(19). Trimmomatic (0.36) (20) was used to remove adapter contamination shown by FastQC. *M.tuberculosis* H37Rv (NC_000962) sequence was used as the reference strain to identify the genetic variations. To avoid errors, four different pipelines were used in data analysis: (a) Maq (Mapping and Assembly with Quality) (0.6.6) (21) (b) Bowtie 2 (2.0.4) was used for variant detection in massively parallel sequencing data (22) in combination with Samtools (23), GATK (3.6-0-g89b7209) and Picard Tools (1.119) (http://broadinstitute.github.io/picard) (24, 25), (c)Varscan 2 (26) and Mpileup of Samtools (23) (d) inGAP, an integrated genome analysis pipeline (27). The consensus of the four software pipeline was considered to call the **S**ingle **N**ucleotide **V**ariations (SNV) in the genome of VPCI591. The variations were mapped using ANNOVAR software (28). We compared VPCI591 with global datasets of variation totaling to 1553, obtained from GMTV (http://mtb.dobzhanskycenter.org) (29) and tbVar database (http://genome.igib.res.in/tbVar/) (30).

### Classification of the genome sequence data

The whole genome SNVs were classified into genic and non-genic/intergenic on the basis of their location on the genome. The genic variations were further classified into (a) Synonymous (SY) (b) Non-synonymous(NS) (c) Stop-gain(SG) (d) Stop-loss(SL) categories depending on their effect on the protein sequence. In case of SG and SL variations we predicted the number of amino acids lost or gained due to premature termination or extension respectively. In SL variations, the extended protein resulting from the addition of amino acids was analysed for its predicted function by BLASTP (31) and InterProScan (32). The ontology for genes harbouring NS variations were carried out using PANTHER (33) and further classified according to the major pathway (4). The non-synonymous SNVs in drug resistance genes were retrieved by literature mining as well as the analysis of TBDReaMDB, a comprehensive drug resistance database (14). The SNVs were screened for their predicted effect through SIFT(34). For non-genic/intergenic variations, the relative distance between the SNV and the downstream gene was determined. From global comparative analysis with tbVar (30) and GMTV (29) database, invariant genes and variations unique to VPCI591 were identified.

### Structural analysis of RpoB (*Rv0667*)

To assess the effect of the NS-SNV in RpoB detected in VPCI591, structural analysis was carried out. The atomic coordinates of RpoB complex with rifampicin (PDB Id: 5UHB; Resolution: 4.29-Å) (35) was obtained from Protein Data Bank (http://www.rcsb.org/pdb). The effect of variation on the structure and the binding of rifampicin was investigated by molecular docking to examine the protein-ligand (rifampicin) interaction using AutoDock 4.2 (36). The structure of the protein with SNVs was generated using SPDBV (37). All the hetero atoms were removed, and both the protein and ligand files were prepared and saved in PDBQT format and used as initial input for AutoDock following the standard protocols. The Lamarckian genetic algorithm (LGA) method was applied for docking and to deal with the protein–antagonist interactions(38). The polar hydrogen atoms were added geometrically and Gasteiger charge to all the atoms of the protein was assigned using AutoDoc tool (ADT). The 3D affinity grid fields with grid map of 50 × 50 × 50 points spaced equally at 0.375 Å and centre of the grid box was 164.1 × 163.34 × 20.42 using auxiliary program AutoGrid to evaluate the binding energies between the ligand and receptor. The standard protocol of ADT utility was used to generate both the grid parameter file (gpf) and docking parameter file (dpf). The resultant docked models of wild-type and mutant complexes with rifampicin were analyzed using Schrodinger and PyMOL to visualize molecular interactions.

### Expression analysis

The intergenic variation mapping between *embC* and *embAB* genes inVPCI591 was validated by Sanger sequencing using the primers, forward (CCTAGGAACGGTGACTCG) and reverse (AGACGACGGCTGCTAGGC). For expression analysis total RNA was extracted from the clinical isolate VPCI591 and H37Rv using the RNeasy mini kit (Qiagen) and was treated with DNase using TURBO™ DNase kit (Invitrogen) and cDNA was prepared using First strand cDNA synthesis kit (Fermentas K1612). Quantitative PCR was performed with *sigA*/*Rv2703* gene as control and fold change was measured by ΔΔCt method using Fast Start universal master mix (Roche). The following primers were used: embA (F-GTAATGAGCGATCTCACCGG/R-CGGTGATCTGGGTGATGTTG); sigA (F-AACGCACCGCCACCAAGTC/R-TGGTGCTGGTCGTAGTGTCCTG).

## Results

### Sequencing of VPCI 591 genome

The sequencing depth of coverage was calculated to be 133X. The FastQC data analysis showed high quality reads after adapter removal. The quality score distribution of all the reads and GC nucleotide distribution of the reads are shown in (Additional File 1: Fig.1A&B). The distribution pattern shows typical pattern of GC rich mycobacterial genomes. The mean quality score distribution is also high (Additional File 1: Fig1C).

**Fig 1:**
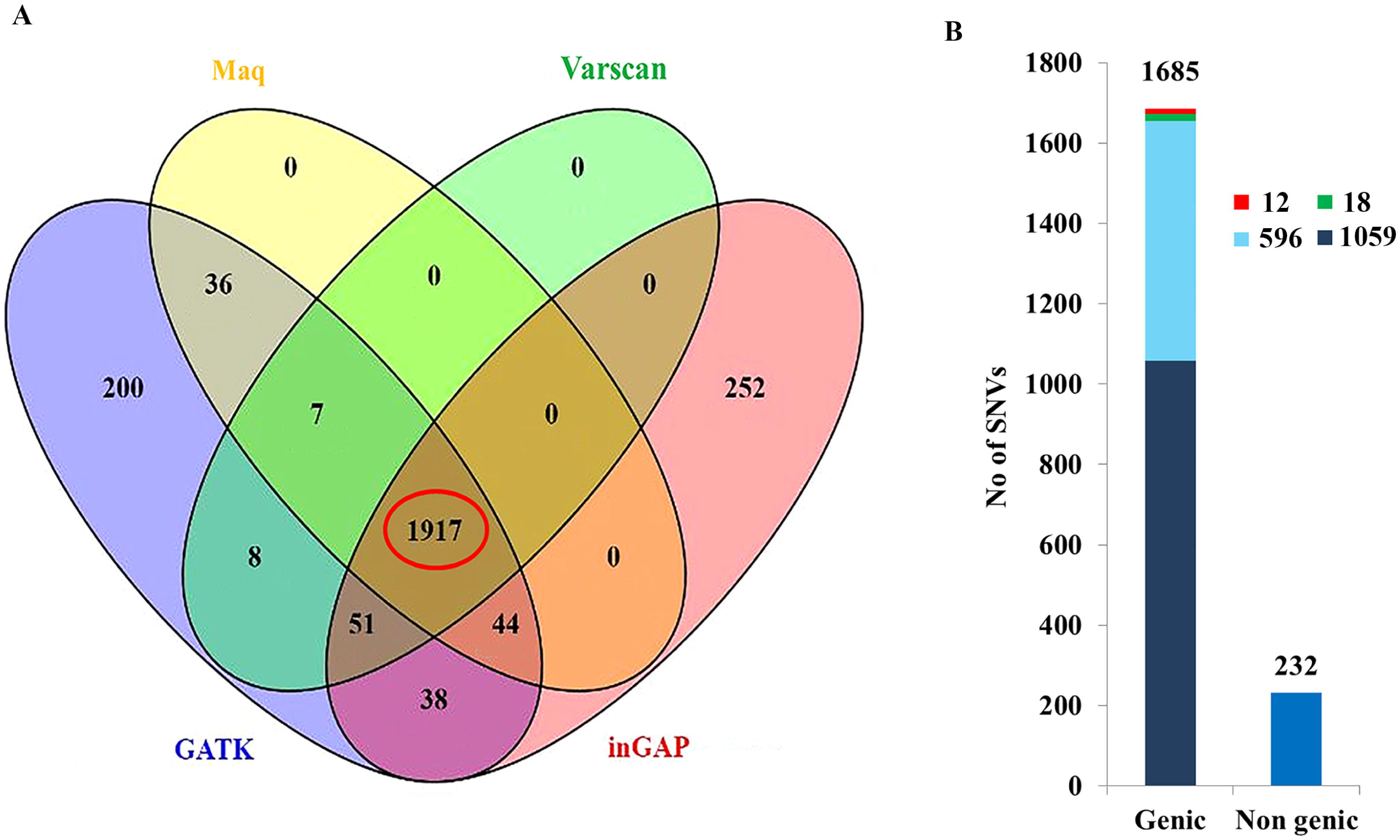
Single Nucleotide variations in VPCI591. **A**. VENN diagram showing the number of SNVs called through the different software and the red circle indicates the consensus. **B.** Total number of variations and their distribution is represented. The Y axis shows the number of variations, the distribution of the 1917 SNVs along with their genic and non-genic distribution and subcategories are shown, stop gain(green), stop-loss(red), synonymous(cyan), non-synonymous(dark blue).

### Analysis of single nucleotide variations; Distribution of genic variations

A total of 1971 SNVs were chosen as consensus variations with high confidence from the four pipelines as described under methods and were considered for further analysis (Fig 1A). 1685 genic and 232 non genic/intergenic variations were obtained. 1059 NS, 596 SY, 18 SG and 12 SL variations were there distributed among 1685 SNVs Fig 1B.

We identified 1685 genic SNVs in VPCI591 distributed among 1245 genes (Fig 2A). We have found 926 genes having single variation among which 737 are NS (Fig 2A,B). The 1059 NS variations found in VPCI591 are distributed in 894 genes (Fig 2B). It is known that the PE-PPE genes are highly variant among clinical isolates (39). In VPCI591, among the PPE genes, *Rv3347c* has 8 SNVs out of which 5 are non-synonymous SNV and *Rv0355c* has 7 SNVs, 5 of which are NS (Fig 2A,B). VPCI591 contains multiple non-synonymous variations which are also seen in 11 genes known for drug resistance and multiple variations are identified in *gyrA, embC, embB* and *rpoB*. Based on SIFT score (<0.05), several variations were predicted to be deleterious; for example A384V in *gyrA*, I21T in *inhA*,N394D in *embC*, P913S in *embA*, D12A in *pncA* and all the 3 variations in *rpoB* (Fig 2C).

**Fig 2:**
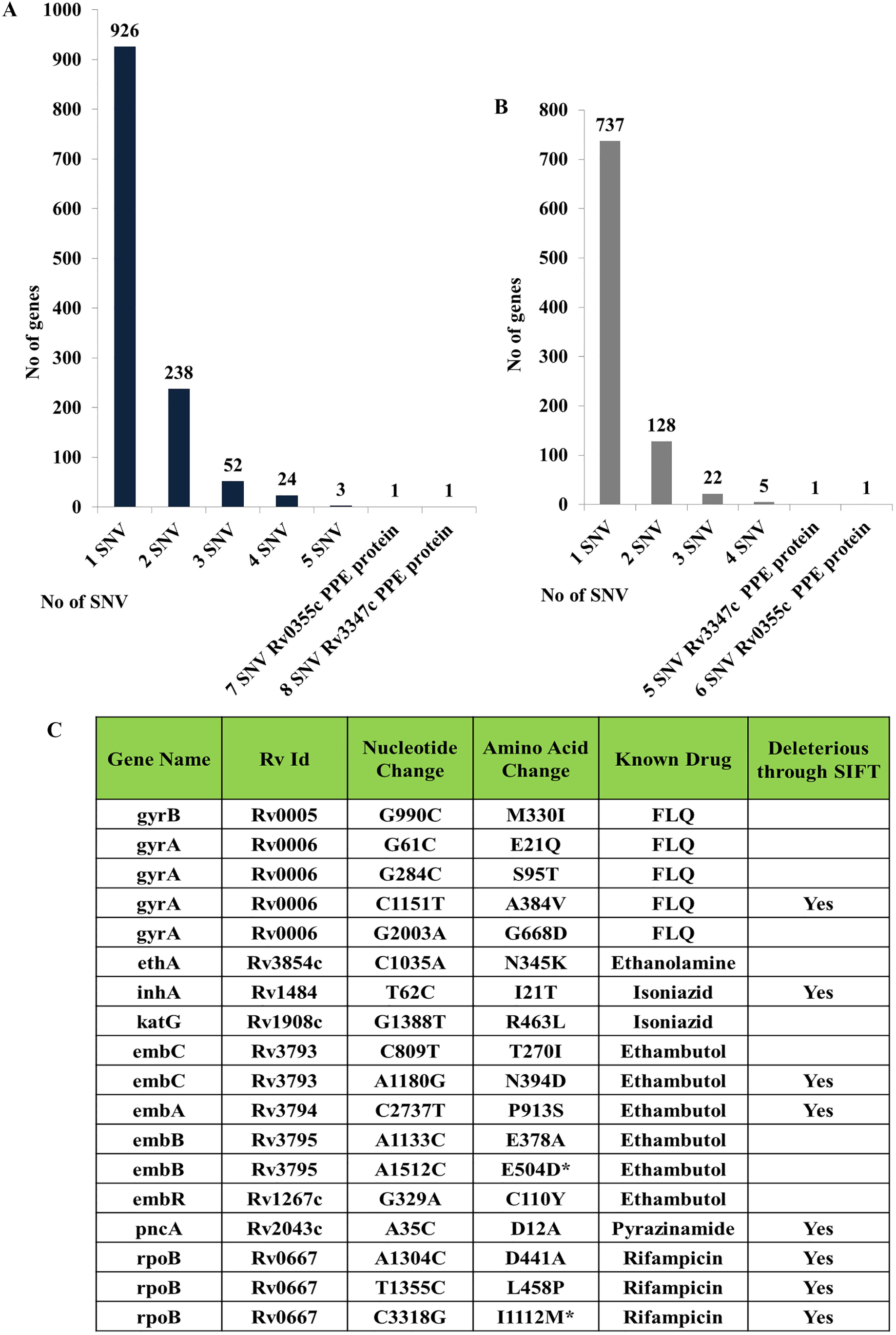
Distribution of Genic SNV in VPCI591. **A.** The overall distribution of genic SNVs. **B.** Distribution non-synonymous (NS) SNVs. **C.** Variation in genes known for drug resistance. FLQ: fluoroquinolones, * Represents variations unique to this clinical isolate.

The genetic variations were found in genes responsible for resistance to other drugs such as, fluoroquinolones, ethanolamine, isoniazid, ethambutol, pyrazinamide and rifampicin (Fig 2C). RNA polymerase β subunit (*rpoB)* harbours 3 variations known to cause resistance to rifampicin (Fig 2C). We found similar variations in 1553 global clinical isolates of *M.tuberculosis*, that we analysed, however the co-occurrence of all the three is unique to VPCI591. In addition, based on SIFT score, we can predict deleterious effect of the SNVs in the following genes;, *mmpL4* (*Rv0450c)* and *mmpL8* (*Rv3823c*) involved in membrane transport, *proC* (*Rv0500*, pyrroline-5-carboxylate reductase), *trcS* (sensor histidine kinase *Rv1032c*), *cyp121*(*Rv2276*, Cytochrome P450 and Polyketide synthases *pks2* (*Rv3825c*)) and *pks7*(*Rv1661*). VPCI591 has SNVs in the two component system genes, *mprA*(*Rv0981*) and *mprB*(*Rv0982*). We have investigated its effect on the expression of immune response genes and also on phago-lysosome fusion (unpublished). A complete list of NS variations with gene names, position of SNV, genomic location and amino acid (AA) change is given in (Additional file 2).

### Functional classification of genes with non-synonymous SNV

The classification of genes containing SNV according to Camus et al., (4) and PANTHER(33) was carried out (Fig 3A). Classification based on Camus et.al., (4) identified 219 genes of intermediary metabolism, respiratory pathway genes, 200 genes involved in cell wall and cell processes, 49 PE and PPE proteins, 65 genes for lipid metabolism of mycobacterium, 45 regulatory proteins and 11 insertion elements,and phage proteins (Fig 3A). A large number of genes (211) are annotated as conserved hypothetical or uncharacterized proteins (Fig 3A).The genes with NS variations (894) were classified according to their molecular function using PANTHER (33). Among the mapped genes, 290 genes are associated with catalytic activity, 56 genes with binding function and 48 genes having transporter activity (Fig 3B). On classifying the catalytic activity genes further, it was observed that 109 of these genes have transferase activity, 88 genes have hydrolase activity and 80 genes have oxidoreductase activity (Fig 3C). Among the 109 genes for transferase activity, 40 genes are involved in transferring acyl group, 31 genes have methyltransferase activity, 12 are glycosyl group transferases and 10 have kinase activity (Fig 3D). The genes in the following classes were under represented among the SNV containing genes: structural constituents of ribosome (GO:0003735), DNA-binding transcription factor activity (GO:0003700), transcription coregulator activity (GO:0003712) and guanyl-nucleotide exchange factor activity (GO:0005085).

**Fig 3:**
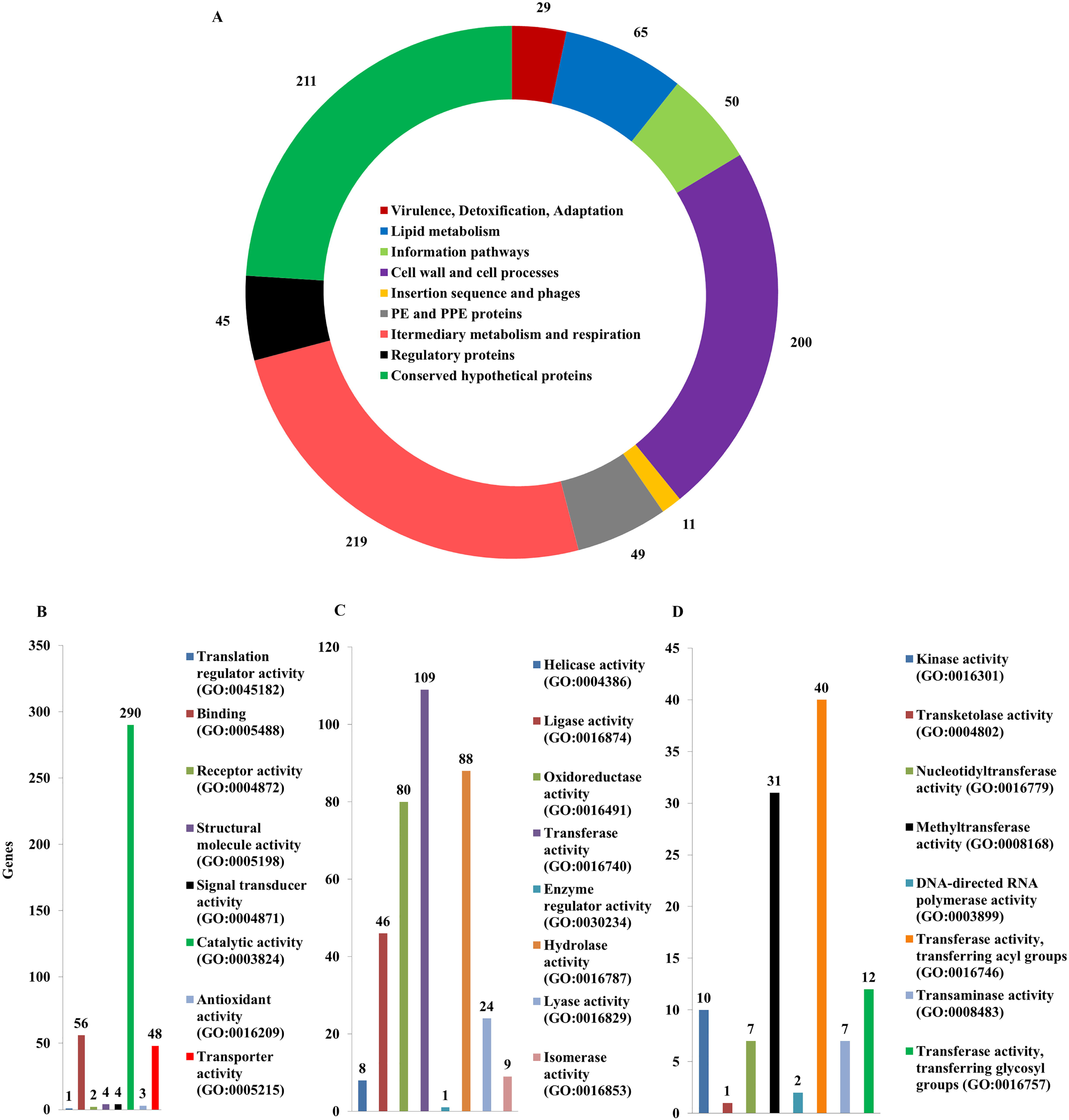
Classification of Genic SNV in VPCI591. **A.** The distribution of non-synonymous SNVs in different functional categories classified according to Camus et al., (2002). **B.** Gene ontological (GO) classification using PANTHER with GO Molecular Function of the genes for NS SNVs. **C.** Sub-classification of catalytic activity highly represented in B, **D**. Sub-classification of transferases that are highly represented in C.

### SNV leading to loss/gain of stop codons

The SNVs that result in a premature stop codon (Stop-Gain,SG) or a loss of stop codon resulting in longer transcripts (Stop-Loss,SL) were identified. The SG variations were identified in 18 genes including ABC transporter *pstA1*(*Rv0930*), dihydroorotate dehydrogenase *pyrD* (*Rv2139*), fatty-acid-CoA synthase *fadD15* (*Rv2187*), NADPH: adrenodoxin oxidoreductase *fprA*(*Rv3106*) and NAD(P)H quinone reductase *lpdA*(*Rv3303c*) (Table 1A). In these cases we identified the domains lost due to which the loss of function is predicted, which was correlated with the loss of a long peptide, including the loss of the functional domain. For example, in *Rv2187* (*fadD15*), 558 amino acids are lost which results in the loss of AMP synthetase/ligase domain. Since many of these genes are annotated as hypothetical proteins, the nature of functional deficiency could not be predicted. Among the 12 stop-loss variations, one of the SNVs results in loss of stop codon at 1467 A>C position in the fatty acid pathway gene polyketide synthase, *pks3* (*Rv1180*) which can now be read through into the adjacent gene, *pks4* (*Rv1181*) to form a longer transcript. This is annotated as *msl3.* Some of the other genes showing stop loss variations are *PPE33*(*Rv1809*), *PPE67*(*Rv3739c*) and epoxide hydrolase e*phF*(*Rv0134*). The predicted new function based on BLAST and InterProScan in VPCI591 due to SL variations are shown with the number of AA added (Table 1B). *Rv0325* gains 155 amino acids to form class I SAM dependent methyl transferase. In *Rv1870*,11 AA were added and it is converted in to an endonuclease.

**Table 1A:** Distribution of stop-gain variation in VPCI591

| **Genomic Position** | **Gene Id** | **Gene Name** | **Nucleotide Change** | **Amino acid changing to stop codon** | **No. AA lost** | **Function** | **Domains lost** |
| --- | --- | --- | --- | --- | --- | --- | --- |
| 162226 | Rv0134 | ephF | G456A | W152X | 148 | Possible epoxide hydrolase EphF | Hydrolase domain |
| 234477 | Rv0197 | Rv0197 | T2247G | Y749X | 14 | Possible oxidoreductase |  |
| 309765 | Rv0257 | Rv0257 | C67T | R23X | 102 | Hypothetical protein |  |
| 549251 | Rv0457c | Rv0457c | G357A | W119X | 555 | Probable peptidase | Oligopeptidase domain |
| 704997 | Rv0610c | Rv0610c | C913T | Q305X | 81 | Hypothetical protein |  |
| 1009490 | Rv0906 | Rv0906 | C547T | Q183X | 190 | Hypothetical protein |  |
| 1037911 | Rv0930 | pstA1 | C913T | R305X | 3 | Probable phosphate-transport integral membrane ABC transporter PstA1 |  |
| 2399782 | Rv2139 | pyrD | C1063T | Q355X | 2 | Probable dihydroorotate dehydrogenase |  |
| 2448288 | Rv2187 | fadD15 | G129A | W43X | 558 | Actelyco asynthetase like | AMP synthetase ligase |
| 2882317 | Rv2563 | Rv2563 | C28T | Q10X | 340 | Probable glutamine-transport transmembrane ABC transporter | Cytoplasmic domain |
| 2929354 | Rv2601 | speE | C967T | Q323X | 202 | Spermidine synthetase |  |
| 2936497 | Rv2608 | PPE42 | C1452A | Y484X | 97 | PPE family protein PPE42 | PPE C-terminal domain |
| 3097349 | Rv2788 | sirR | C391T | Q131X | 130 | Probable transcriptional repressor SirR |  |
| 3351472 | Rv2994 | Rv2994 | G204A | W68X | 377 | Probable integral membrane protein |  |
| 3378720 | Rv3019<br>c | esxR | G282A | W94X | 3 | Secreted ESAT-6<br>like protein<br>EsxR |  |
| 3442240 | Rv3079<br>c | Rv3079c | G358T | E120X | 156 | Hypothetical<br>protein | Luciferase<br>like<br>domain |
| 3689523 | Rv3303<br>c | lpdA | C1416A | C472X | 22 | NAD(P)H<br>quinone<br>reductase LpdA |  |
| 4351039 | Rv3872 | PE35 | G295T | E99X | 1 | PE family-<br>related protein<br>PE35 |  |

**Table 1B:**
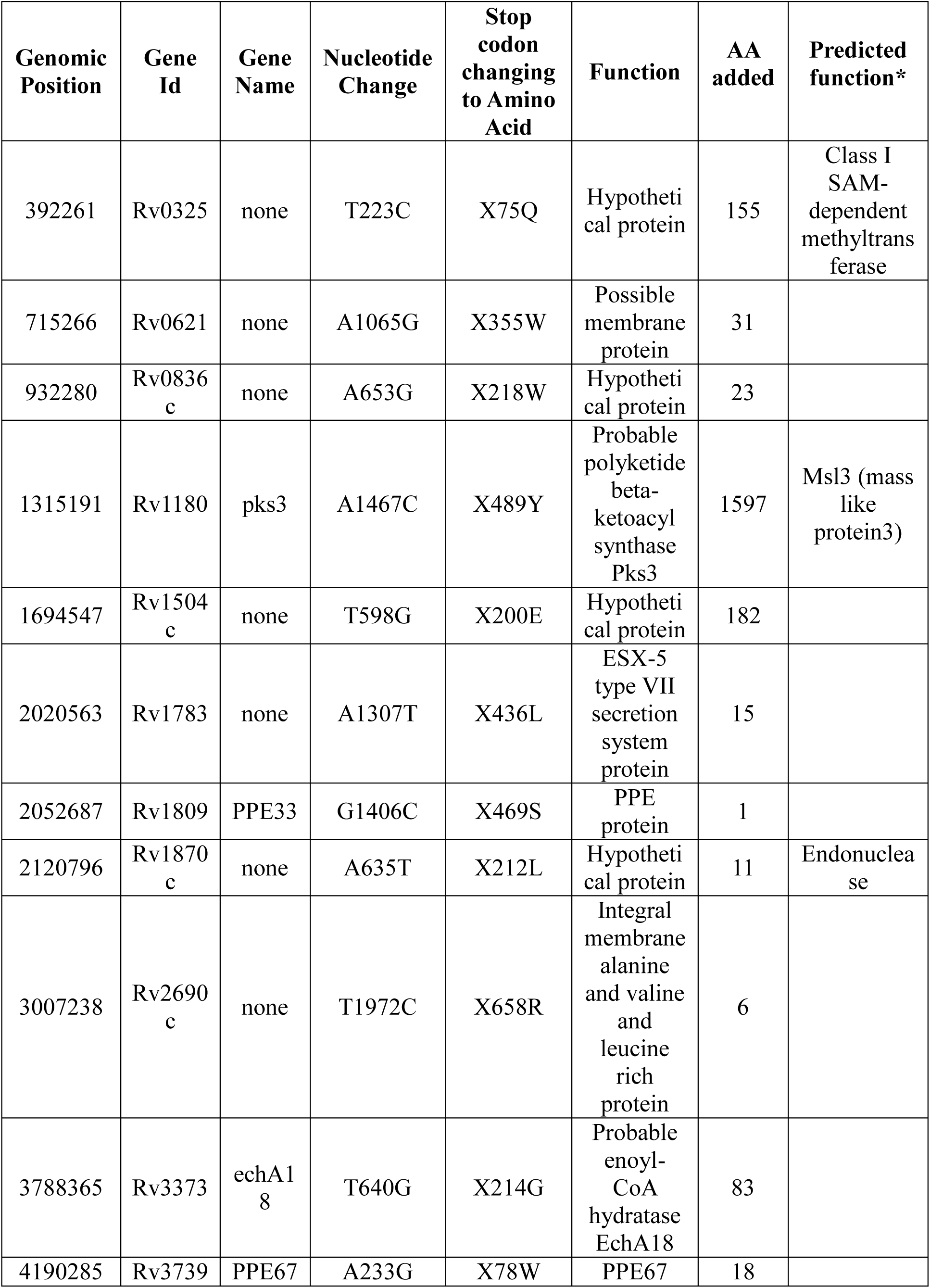
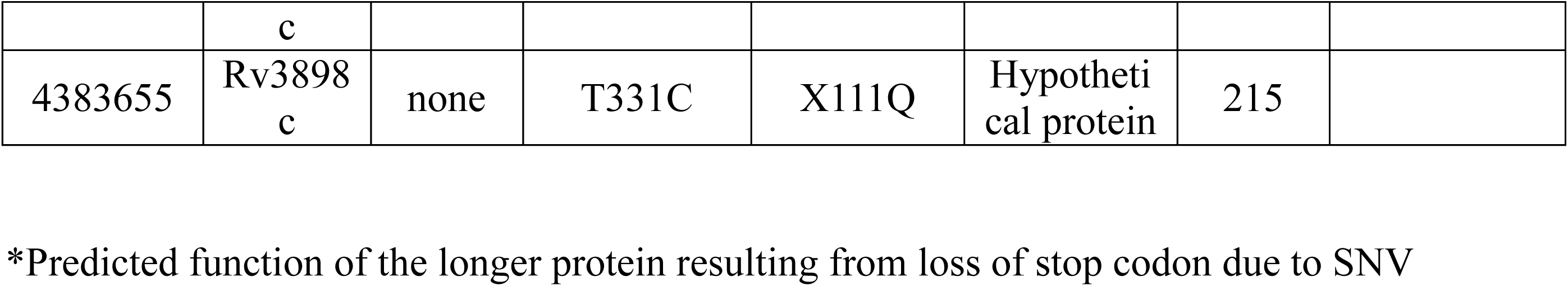
Distribution of stop-loss variation in VPCI591

### Variation in intergenic region between divergently transcribed genes

There are several intergenic regions that are flanked by divergently transcribed genes. The presence of promoters or regulatory regions within such intergenic non-coding regions are known in *M.tuberculosis* (40). Thus intergenic SNVs can affect the transcription/regulation of the downstream gene. We have mapped the intergenic variations occurring upstream of genes and also those mapping between divergently transcribed genes (Fig 4A; I and II) and computed the relative distance between the SNV position and the gene(s). We have identified a total of 232 intergenic variations, 32 of them are positioned between genes transcribing in opposite directions and these have the potential to affect both the genes. These were binned into categories depending upon the distance between the downstream genes (Fig 4B). There are 77 variations that map within 50 bp from transcription start site, 39 SNVs between 50-100bp region and 35 SNVs between 100-150bp regions indicating that majority of the intergenic variations are close to the transcription start site. Such variations in putative promoter or regulatory variations can affect both the flanking genes transcribed in opposite directions. Some of the genes whose transcription could be affected by the SNV in VPCI591 were identified (Fig 4C). The complete list of the intergenic variations is given in (Additional file 3 and Additional file 4) respectively. We found a variation within the 85 bp intergenic region between *embC* and *embAB*, 4 bp upstream of transcription start site of *embA.* The evaluation of the functional significance of this variation is discussed later.

**Fig 4:**
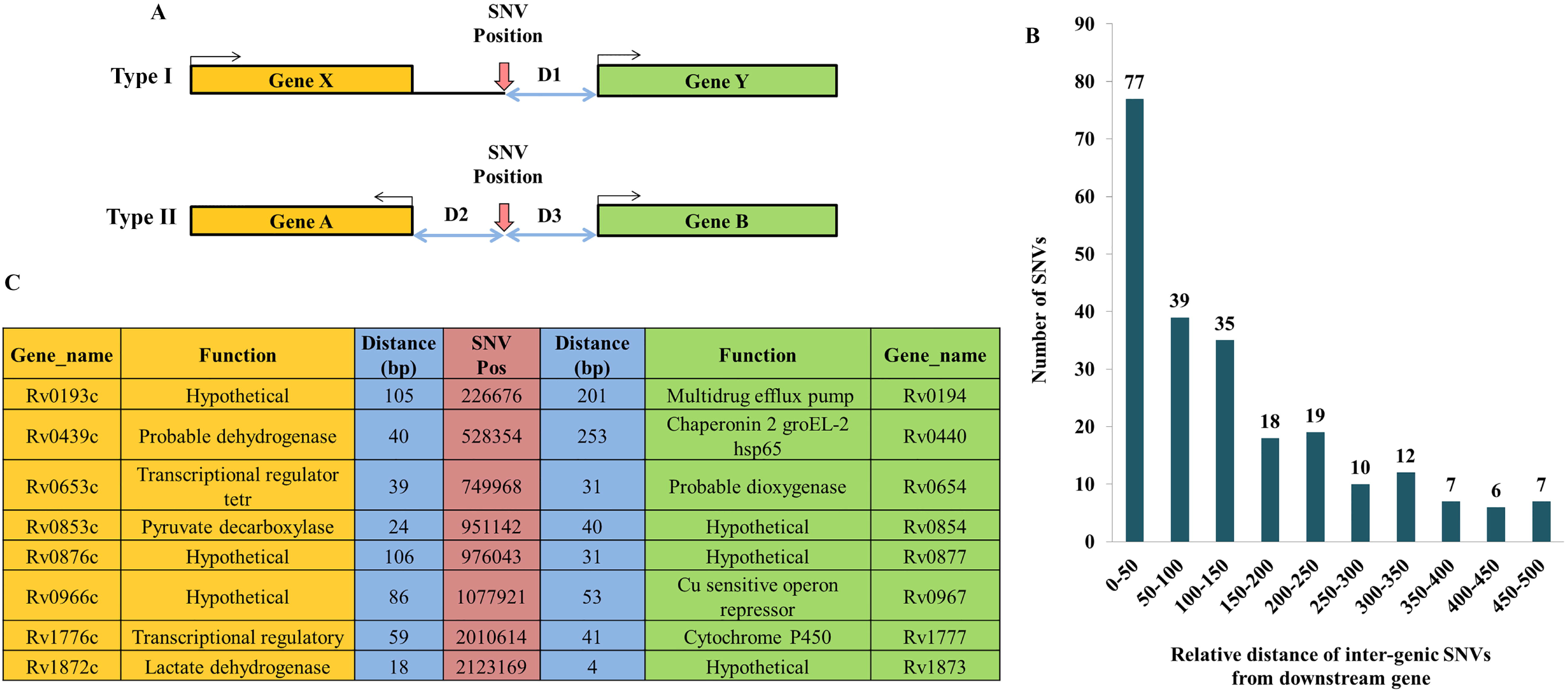
Intergenic SNV in VPCI591. **A**. Schematic representation of intergenic variations leading to type I and type II configuration with reference to the direction of transcription of the adjacent genes. SNV position is indicated with vertical red arrows. The two-way arrows (D1,D2,D3), show the distance between the SNV and the downstream genes, bent arrows indicate direction of transcription. **B.** Distribution of non-genic SNVs relative to the distance from the downstream gene (Type I and II) **C.** The list flanking genes and their relative distance (D2 and D3) from the intergenic SNVs in Type II configuration.

### Unique variations in VPCI591 and invariant genes in Clinical isolates of *M.tuberculosis*

We found several invariant genes among the 1553 clinical isolates and also VPCI591. This includes genes coding for secretory proteins (*Rv3875*/ESAT-6), EsxM), ribosomal proteins (RpmJ and RpmH, RpsJ, RpsH), toxin-antitoxin pathway proteins (VapB3, VapB26, VapC7, VapB10), several transposase for IS elements (Rv2278, Rv2354, Rv2355) and many putative PhiRv2 prophage proteins (Rv2654c, Rv2656c, Rv2657c) known in *M.tuberculosis.* Interestingly there are several PE, PPE and PGRS genes identified like *Rv2099c* (*PE21*), *Rv1195* (*PE13*), *Rv2431c* (*PE25*), *Rv3622c* (*PE32*), *Rv2098c* (PGRS36) proteins that do not vary among the genomes we compared (1553). The complete list of the invariant genes is shown in (Additional file 5).

The comparative analysis with global clinical isolates, led to the identification of 143 unique variations in VPCI591, among which 125 are genic and 16 are intergenic. Most of the genic variations were non-synonymous and stop-gain SNVs. Ontology classification revealed that majority of them fall under genes with catalytic function (75%) and transportation (17%). The complete list of the unique genes with amino acid alteration along with the SNV position in the genome is shown in (Additional file 6). A number of intergenic variations lie within the known promoter/regulatory regions. Intergenic variation in the cis-regulatory sequences of mce1 operon between *Rv0166* (fadD5) and *Rv0167*(yrbE1A) were identified(40). SNV was identified within IS 1560 elements and mutation hotspot for anti-tuberculosis drug ethambutol, between *embC* and *embAB* gene. The complete list of unique intergenic variations identified in VPCI591 along with their genomic position and the adjacent genes is shown in (Additional file 7)

### The effect of SNV in RpoB subunit and its possible implication

As mentioned earlier, 3 variations co-occurring in RpoB (*Rv0667*) at D441A, L458P and I1112M which is unique to VPCI591. We examined the effect of these on the structure of RpoB and its interaction with rifampicin. To understand the binding of rifampicin at the active site of RpoB, molecular docking analysis was performed using the crystal structure of RpoB (PDB Id: 5UHB). The locations of the 3 variations are mapped in the crystal structure at D441, L458 and I1112 (Additional file 8: Fig A). The wild type protein showed His451, Phe439, Gln438, Arg454, Arg465, Ser456, Asn493 interacting with the drug rifampicin at the active site (Fig 5A,C). Each variation was examined for its effect separately as well as in combination with each other; D441A showed loss of all the interactions with drug except Gly438, Phe439 and Arg465 (Additional file 8: Fig B&C)and, while L458P also showed loss of all interactions with drug except Gln438, His441, Phe439 and Arg465 (Additional file 8: Fig D and E). The variation I1112M showed loss of interactions with drug except Gln438, Phe439 and Arg465 (Additional file 8: Fig F&G). The combination of all the three variations, D441A, L458P and I1112M as present in the clinical isolate VPCI591, showed loss of all interactions with the drug retaining the interaction with only with Arg454 (Fig 5B&D). The binding energy and the respective Ki values for the 3 variations were calculated for individual variations. The wild type shows (ΔG) −9.27 kcal/mol while the variants D441A, L458P and I1112M have −6.67 kcal/mol, −7.11 kcal/mol and −7.18 kcal/mol respectively. The combination of the three, D441A+L458P+I1112M has ΔG equal to −5.82 kcal/mol showing loss of binding of rifampicin at the active site. This could result in drug resistance. The Ki values calculated from the wild type is 160nM whereas RpoB from VPCI591 shows 53.96μM. The complete representation of the individual variations as well as the combined, the number of hydrogen bonding and the relative distances in Å is shown in (Table 2).

**Table 2.** Molecular docking analysis of RpoB crystal structure of H37Rv PDB-5UHB with Rifampicin by Autodock 4.2.

| Protein Model | Binding Energy, $\Delta G$ (Kcal mol<sup>-1</sup>) | $K_i$ observed (nM/μM) | Hydrogen Bonding | |
| --- | --- | --- | --- | --- |
|  |  |  | Residues | Distance (Å) |
| H37Rv | -9.27 | 160.26 (nM) | Gln438 | 2.9 |
|  |  |  | Phe439 | 2.9, 3.1 |
|  |  |  | His451 | 3.2 |
|  |  |  | Arg454 | 2.8 |
|  |  |  | Ser456 | 2.8 |
|  |  |  | Arg465 | 3.5 |
|  |  |  | Asn493 | 3.2 |
| D441A* | -6.76 | 11.06 (μM) | Gln438 | 3.2 |
|  |  |  | Phe439 | 2.7, 2.7 |
|  |  |  | Arg465 | 3.4 |
| L458P* | -7.11 | 6.16 (μM) | Gln438 | 2.8 |
|  |  |  | Phe439 | 2.7, 2.8 |
|  |  |  | His441 | 3.2 |
|  |  |  | Arg465 | 3.3, 3.5 |
| I1112M* | -7.18 | 5.50 (μM) | Gln438 | 2.8 |
|  |  |  | Phe439 | 2.6, 2.8, 3.3 |
|  |  |  | Arg465 | 3.5 |
| D441A+L458P+I1112M** | -5.82 | 53.96 (μM) | Arg454 | 3.1, 3.1 |

Binding parameters computed based on molecular docking analysis of the crystal structure of RpoB H37Rv (PDB-5UHB) with Rifampicin by Autodock 4.2. The binding parameters were calculated for RpoB of H37Rv and VPCI591; \* with individual SNV and \*\* the three SNVs co-occurring in VPCI591.

**Fig 5:**
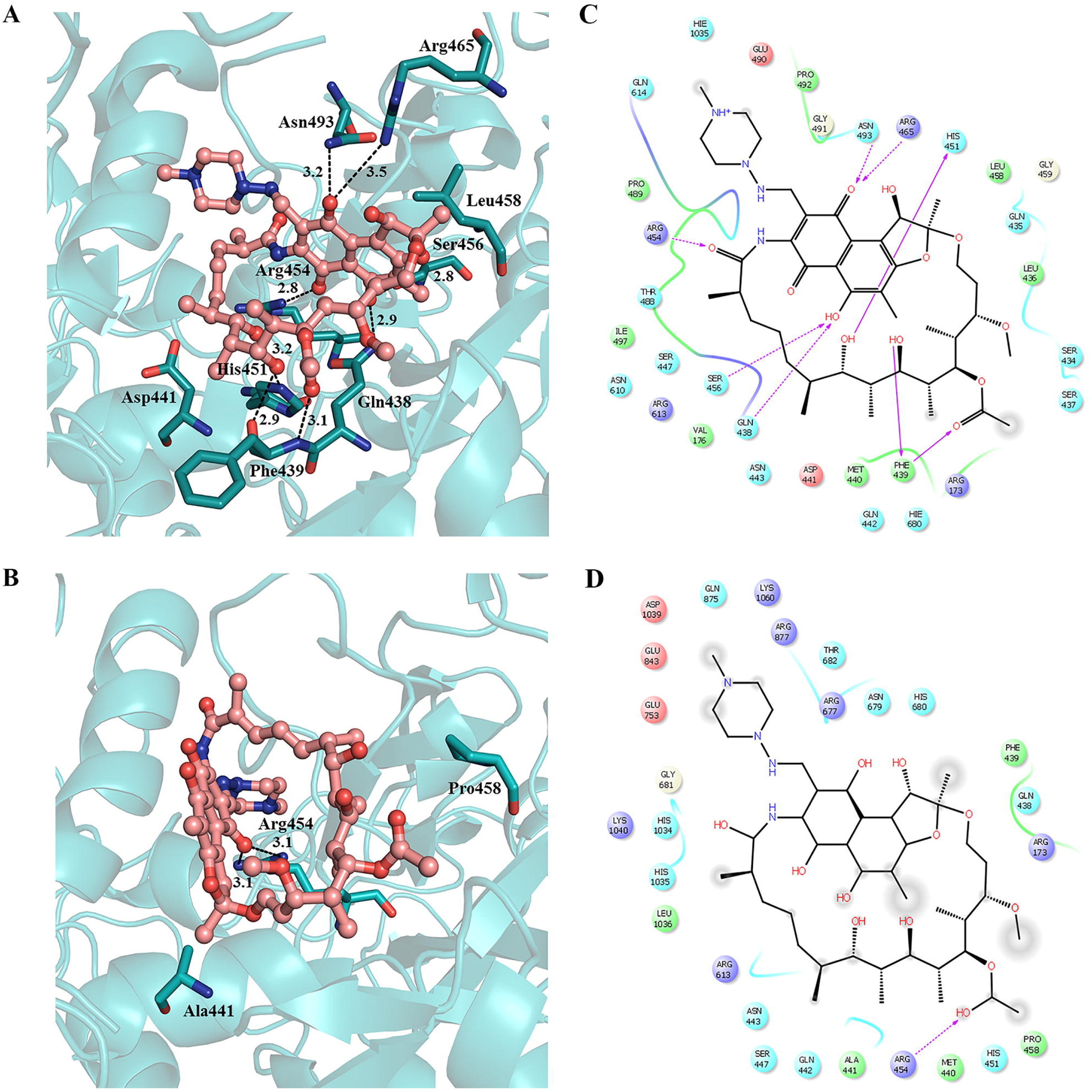
Docking analysis of SNV in RpoB. A & C are for (RpoB-Wild type-H37Rv) and B & D are for (RpoB-clinical isolate-VPCI591). A & B represent 3D view, amino acid residues are shown with sticks and rifampicin (red) is shown as ball and stick model, hydrogen bonds as dotted lines (black). C & D represent the corresponding 2D sketch of molecular interaction of rifampicin with adjacent amino acid residues indicated by purple arrow.

### Effect of SNV in intergenic region upstream of *embA* (*Rv3794*) gene

Several NS variations were identified with the associated genes for ethambutol resistance in *embC* (C809T, A1180G), *embA* (P913S), *embB* (E378A, E504D) (Fig 6A). We found NS variation at *embR* (C110Y) which is not a part of the operon. In addition to the genic variations, an intergenic SNV at position 4243299 C>T at −4bp position upstream to *embA* gene is identified in VPCI591. The SNV was confirmed by PCR followed by Sanger sequencing. The expression level of *embAB* was compared with that of the reference strain H37Rv by quantitative PCR, using expression of *sigA* for normalization. No SNV is identified in *sigA* in VPCI591. There is 3.6 fold increases in the expression of *embA* gene in VPCI591 as compared to H37Rv in log phase and 1.25 fold increase in expression in the stationary phase of growth, indicating a gain-of-function due to the intergenic variation (Fig 6B).

**Fig 6:**
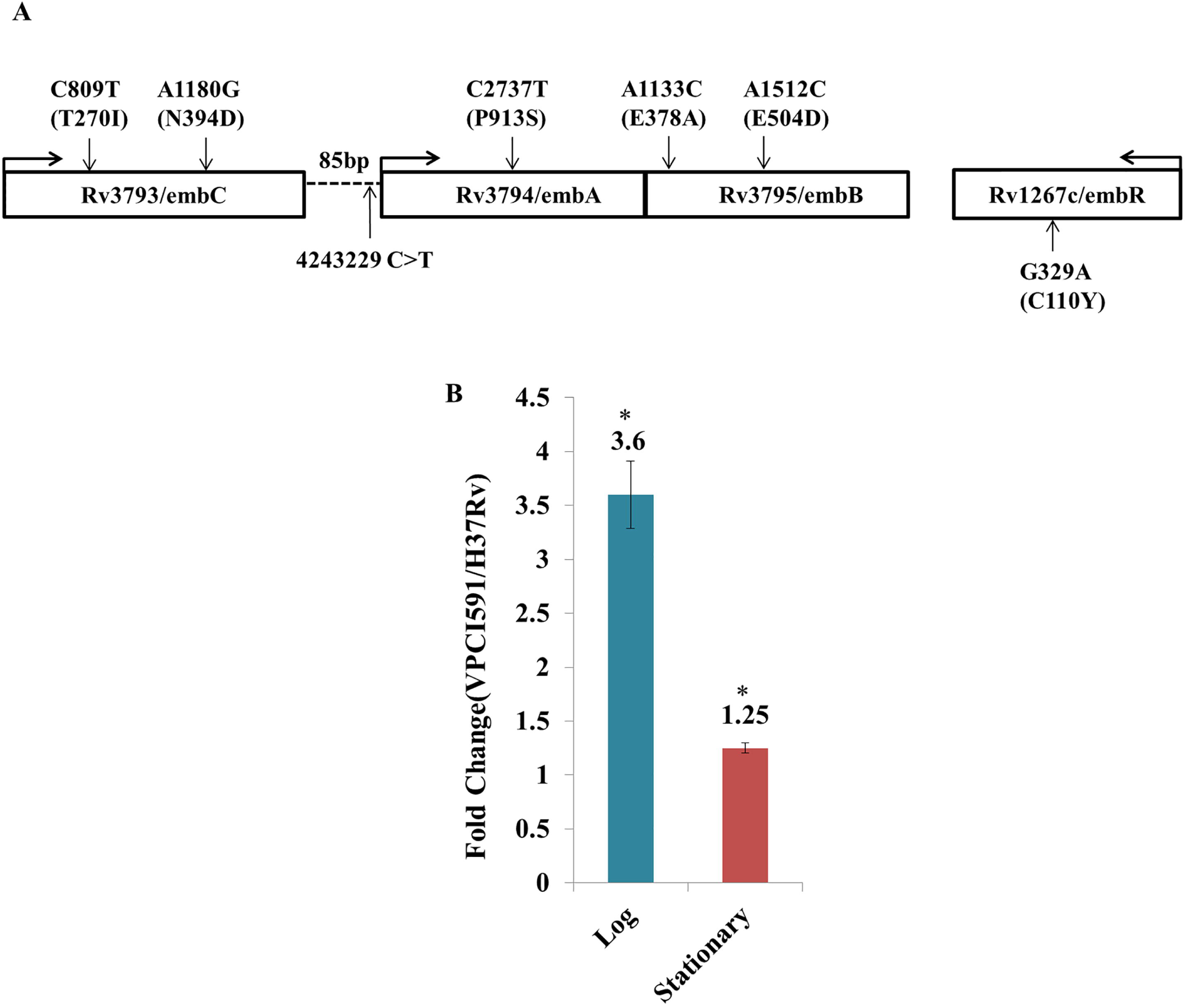
Effect of intergenic SNV on the expression of *embAB*. **A.** Diagrammatic representation of *embCAB* operon of VPCI591 with the SNVs (vertical arrows). The direction of transcription is indicated by horizontal arrows. The intergenic region between *embC* and *embAB* (85 bp) is represented by dotted lines. **B.** Relative expression of *embAB* gene in VPCI591 with respect to H37Rv at log phase and stationary phase, represented as mean ± SEM (*p<0.05). The results from two biological replicates are shown.

## Discussion

The directed polymorphism analysis of the *mce* 1 operons in VPCI591, led to the identification of gain of function SNV at 196800 (G>C) in the intergenic region between *Rv0166-Rv0167*(40). In the light of this, the whole genome sequencing and analysis was taken up to map the non-synonymous SNVs in VPCI591. This led us to predict the molecular correlates for the resistance phenotype of VPCI591 for two of the first line drugs, rifampicin and ethambutol. VPCI591 is resistant to significantly higher concentration of most of the first-line drugs compared to single/two drug resistance strains (17). Here we find that the co-occurrence of three genic non-synonymous SNVs which is unique to VPCI, weakens the interaction of RNA polymerase β subunit with rifampicin, as reflected in the increase in binding affinity from −9.27 kcal/mol to −5.82 kcal/mol as well as increase in Ki values from 160.26 (nM) to 53.96 (μM). However the effect of SNVs on the affinity of RpoB for rifampicin as assessed by the structural perspective alone does not entirely justify this level of resistance. It is known that in addition to *RpoB* mutation, the active efflux pumps like *Rv1258c, Rv1410c*, and *Rv0783*, the major facilitator superfamily (MFS) proteins, contribute to rifampicin resistance (41). VPCI591 bears no variation in these genes except SNV in *Rv2936*/*drrA*(H309D), implicated in rifampicin efflux (41). VPCI591 has NS variations in *rpoC* (A172V) and no variation in *rpoA*. The presence of these off-target variations are also known to contribute to the evolution and survival of drug-resistant *M. tuberculosis* (42, 43).

Among the high confidence SNVs identified, the antigenic *PE-PPE* proteins show high variation, having multiple variations in a given gene in VPCI591 as in the 1553 global clinical isolates of *M.tuberculosis*, that we analysed from the repository tbVar (30) and GMTV (29). However there are certain invariant *PE, PPE* family of genes. The number of SNVs per gene is highest in the PPE genes *Rv0355c* and *Rv3347c*; out of the 7 SNVs in *Rv0355c*, 6 are non-synonymous, while 5 out of 8 SNVs in *Rv3347c* are non-synonymous. The members such as *PPE 35, PPE55, PPE8, PPE54, PPE34, PPE24* genes are known to be highly polymorphic among clinical isolates (39).SNVs leading to stop-loss (SL) in *PPE33, PPE67* and stop-gain(SG) in *PPE42, PPE35*, in addition to SG in fatty acid ligase gene, *fadD15*. However the phenotypic consequence of these variations is not reflected in the ability of the pathogen to infect and survive in the human host or in in-vitro growth.

Apart from the genic variations, the intergenic variations can have significant effect on the expression of the genes if they map within putative promoters or regulatory regions. The SNVs mapping in the intergenic region between two genes that are transcribed in opposite directions, can impact the transcription of both the genes. There are several such instances in VPCI591, including SNV at 2010614, between *Rv1776c* (transcriptional regulator) and *Rv1777* (Cytochrome P450), *Rv2333c* (Drug efflux protein) and *Rv2334* (Cysteine synthase).

It is known that 50% to 70% of drug resistant clinical isolates of *M.tuberculosis* have ethambutol resistance and carry a mutation in a relatively short region in *Rv3795*/*embB* gene, primarily at codon positions 306, 406 and 497. They represent promising diagnostic markers for the rapid detection of EMB resistant tuberculosis strains (15, 44-47). Out of the various mechanisms of ethambutol resistance in *M.tuberculosis*, the over expression of *embAB* gene due to the presence of intergenic mutation, is one of the most well-understood mechanism (48).

VPCI591 harbours several SNVs which might be associated with ethambutol resistance; in *embC*/*Rv3793* (T270I) and (N394D), *embA*/*Rv3794*(P913S) and two variations in *embB*/*Rv3795* (E378A) and (E504D). There is also another non-synonymous change in *embR*/*Rv1267c* (C110Y). In addition, VPCI591 has a variation in *pknH*/*Rv1266c*(R607Q). *PknH*-mediated increase in the transcription of *embAB* genes significantly alters resistance to ethambutol (49). Thus the SNVs in several genes conferring resistance to ethambutol, and a threefold increase in the transcription of *embAB* collectively contribute to the high MIC for ethambutol, 15μg/ml of VPCI591 (17) relative to MIC of 0.5-2 μg/ml of H37Rv (17, 50).

The genes involved in transferase activity were the major functional class represented among the non-synonymous SNV containing genes in VPCI591 as also in others as reflected in tbVar(30) and GMTV(29) databases. It can be speculated that transferases including those involved in the post-translational modifications can tolerate variations and thus plasticity, due their important role in conferring functional diversity.

The identification of genes that do not show any SNV in a large number of global clinical isolates (invariant regions), would be potential diagnostic markers for identification of *M.tuberculosis* complex (51-53). We found several invariant genes in 1553 global clinical isolate including VPCI591. It has been shown that the invariant region in 190bp region of *Rv1458c* and partial regions of *Rv0440* can be used as diagnostic markers for identification of *M.tuberculosis* complex (MTBC)(51). The major group of invariant genes in our study includes, toxin antitoxin pathway genes like *Rv0661c, Rv2760c, Rv0664*, and *Rv0581*, and transposase of IS elements (Insertion elements) like *IS6110*. Several ribosomal subunit genes and the putative mycobacteriophage proteins like probable PhiRv1 phage protein are also devoid of variations in the clinical isolates. Several secretory pathway proteins like *Rv3875* (ESAT-6), TatA, SecG and sugar transport proteins like SugB, proteins of electron transport chain *nuoE, nuoC, nuoK* are also found invariant. Thus such invariant regions present in clinical isolates can be used as diagnostic markers.

## Conclusion

The genetic variation in *M.tuberculosis* clinical isolates is common among almost all functional classes of genes. We demonstrate that the variations bring about quantitative changes in transcription and can lead to altered structure of protein and interaction with the drug molecule. Likewise intergenic distance for non-genic SNVs plays a very important role if they lie within the putative promoter or regulatory region. However, the clinical isolates survive and are pathogenic implying that there is no drastic negative effect that is obvious. On the other hand, the invariant genes can be explored for their potential as drug targets.

## Declarations

### Availability of data and materials

NCBI: SRX5802345, Bioproject accession number PRJNA540936, BioSample accession numbers: SAMN11568242.

Additional files 1-8

### Ethics approval and consent to participate

Not applicable

### Consent for publication

All the authors consented for publication

## Acknowledgement

The authors acknowledge Dr. Vinod Scaria at CSIR-Institute of Genomics and Integrative Biology (IGIB) for his help with the genome sequencing.

## Authors’ contributions

M.B., V.B., K.B., M.V. conceived and designed the experiments. K.B., V.N., and M.J., performed the experiments. K.B., V.N. VB analysed the data and wrote the manuscript. R.P provided inputs through discussion. All authors discussed and read the manuscript.

---

**Additional file 1: FastQC analysis. A.** Assessment of quality of the sequence reads. The data shows quartile deviation with mean deviation (blue curve). Sequence quality is shown on the X axis and base position on Y axis. Green-high quality reads, yellow-average and red-low quality reads. **B.** Mean distribution of GC content, estimated from VPCI591 sequence data (red) compared to the theoretical distribution of the GC rich genome (blue). **C**. Mean sequence quality score, ranging from 32-34 indicating high quality reads.

**Additional file 8: Docking analysis of SNV in RpoB**. A. The crystal structure of RpoB (5UHB) is represented harboring all the 3 variations, as present in VPCI591 in green. B,D and F represents the 3D view for D441A, L452P & Ill12M respectively. Amino acid residues are shown with sticks and rifampicin (red) is shown with ball and stick model. Hydrogen bonds are shown as broken line (black). C, E and G represent the corresponding 2D sketch. Rifampicin is represented in black interacting to amino acids (color circles) through purple arrows.

## References

1. Cubillos-Ruiz A, Morales J, Zambrano MM. Analysis of the genetic variation in Mycobacterium tuberculosis strains by multiple genome alignments. BMC research notes. 2008;1(1):110.

2. Cole S, Brosch R, Parkhill J, Garnier T, Churcher C, Harris D, et al. Deciphering the biology of Mycobacterium tuberculosis from the complete genome sequence. Nature. 1998;393(6685):537.

3. Behr M, Wilson M, Gill W, Salamon H, Schoolnik G, Rane S, et al. Comparative genomics of BCG vaccines by whole-genome DNA microarray. Science. 1999;284(5419):1520–3.

4. Camus J-C, Pryor MJ, Médigue C, Cole ST. Re-annotation of the genome sequence of Mycobacterium tuberculosis H37Rv. Microbiology. 2002;148(10):2967–73.

5. Advani J, Verma R, Chatterjee O, Pachouri PK, Upadhyay P, Singh R, et al. Whole Genome Sequencing of Mycobacterium tuberculosis Clinical Isolates From India Reveals Genetic Heterogeneity and Region-Specific Variations That Might Affect Drug Susceptibility. Frontiers in microbiology. 2019;10:309.

6. Brown AC, Bryant JM, Einer-Jensen K, Holdstock J, Houniet DT, Chan JZ, et al. Rapid Whole-Genome Sequencing of Mycobacterium tuberculosis Isolates Directly from Clinical Samples. J Clin Microbiol. 2015;53(7):2230–7.

7. Kidenya BR, Mshana SE, Fitzgerald DW, Ocheretina O. Genotypic drug resistance using whole-genome sequencing of Mycobacterium tuberculosis clinical isolates from North-western Tanzania. Tuberculosis (Edinb). 2018;109:97–101.

8. Takiff HE, Feo O. Clinical value of whole-genome sequencing of Mycobacterium tuberculosis. The Lancet Infectious Diseases. 2015;15(9):1077–90.

9. Roa MB, Tablizo FA, Morado EKD, Cunanan LF, Uy IDC, Ng KCS, et al. Whole-genome sequencing and single nucleotide polymorphisms in multidrug-resistant clinical isolates of Mycobacterium tuberculosis from the Philippines. Journal of global antimicrobial resistance. 2018;15:239–45.

10. Ford C, Yusim K, Ioerger T, Feng S, Chase M, Greene M, et al. Mycobacterium tuberculosis–heterogeneity revealed through whole genome sequencing. Tuberculosis. 2012;92(3):194–201.

11. Gagneux S, Small PM. Global phylogeography of Mycobacterium tuberculosis and implications for tuberculosis product development. The Lancet Infectious Diseases. 2007;7(5):328–37.

12. Gao Q, Kripke KE, Saldanha AJ, Yan W, Holmes S, Small PM. Gene expression diversity among Mycobacterium tuberculosis clinical isolates. Microbiology. 2005;151(1):5–14.

13. Rehren G, Walters S, Fontan P, Smith I, Zárraga AM. Differential gene expression between Mycobacterium bovis and Mycobacterium tuberculosis. Tuberculosis. 2007;87(4):347–59.

14. Sandgren A, Strong M, Muthukrishnan P, Weiner BK, Church GM, Murray MB. Tuberculosis drug resistance mutation database. PLoS medicine. 2009;6(2):e1000002.

15. Zhang H, Li D, Zhao L, Fleming J, Lin N, Wang T, et al. Genome sequencing of 161 Mycobacterium tuberculosis isolates from China identifies genes and intergenic regions associated with drug resistance. Nature genetics. 2013;45(10):1255.

16. Sharma M, Bose M, Sharma L, Diwakar A, Kumar S, Gaur SN, et al. Intracellular survival of Mycobacterium tuberculosis in macrophages is modulated by phenotype of the pathogen and immune status of the host. International journal of mycobacteriology. 2012;1(2):65–74.

17. Tandon R, Ponnan P, Aggarwal N, Pathak R, Baghel AS, Gupta G, et al. Characterization of 7-amino-4-methylcoumarin as an effective antitubercular agent: structure–activity relationships. Journal of antimicrobial chemotherapy. 2011;66(11):2543–55.

18. Bose M, Chander A, Das R. A rapid and gentle method for the isolation of genomic DNA from mycobacteria. Nucleic acids research. 1993;21(10):2529.

19. Andrews S. FastQC: a quality control tool for high throughput sequence data. 2010.

20. Bolger AM, Lohse M, Usadel B. Trimmomatic: a flexible trimmer for Illumina sequence data. Bioinformatics. 2014;30(15):2114–20.

21. Li H, Ruan J, Durbin R. Maq: Mapping and assembly with qualities. Version 06. 2008;3.

22. Li H, Durbin R. Fast and accurate short read alignment with Burrows–Wheeler transform. bioinformatics. 2009;25(14):1754–60.

23. Li H, Handsaker B, Wysoker A, Fennell T, Ruan J, Homer N, et al. The sequence alignment/map format and SAMtools. Bioinformatics. 2009;25(16):2078–9.

24. McKenna A, Hanna M, Banks E, Sivachenko A, Cibulskis K, Kernytsky A, et al. The Genome Analysis Toolkit: a MapReduce framework for analyzing next-generation DNA sequencing data. Genome research. 2010.

25. Van der Auwera GA, Carneiro MO, Hartl C, Poplin R, Del Angel G, Levy-Moonshine A, et al. From FastQ data to high-confidence variant calls: the genome analysis toolkit best practices pipeline. Current protocols in bioinformatics. 2013;43(1):11.0. 1-.0. 33.

26. Koboldt DC, Zhang Q, Larson DE, Shen D, McLellan MD, Lin L, et al. VarScan 2: somatic mutation and copy number alteration discovery in cancer by exome sequencing. Genome research. 2012.

27. Qi J, Zhao F, Buboltz A, Schuster SC. inGAP: an integrated next-generation genome analysis pipeline. Bioinformatics. 2009;26(1):127–9.

28. Wang K, Li M, Hakonarson H. ANNOVAR: functional annotation of genetic variants from high-throughput sequencing data. Nucleic acids research. 2010;38(16):e164–e.

29. Chernyaeva EN, Shulgina MV, Rotkevich MS, Dobrynin PV, Simonov SA, Shitikov EA, et al. Genome-wide Mycobacterium tuberculosis variation (GMTV) database: a new tool for integrating sequence variations and epidemiology. BMC Genomics. 2014;15:308.

30. Joshi KR, Dhiman H, Scaria V. tbvar: a comprehensive genome variation resource for Mycobacterium tuberculosis. Database. 2014;2014.

31. Altschul SF, Gish W, Miller W, Myers EW, Lipman DJ. Basic local alignment search tool. Journal of molecular biology. 1990;215(3):403–10.

32. Zdobnov EM, Apweiler R. InterProScan–an integration platform for the signature-recognition methods in InterPro. Bioinformatics. 2001;17(9):847–8.

33. Mi H, Muruganujan A, Casagrande JT, Thomas PD. Large-scale gene function analysis with the PANTHER classification system. Nature protocols. 2013;8(8):1551.

34. Ng PC, Henikoff S. SIFT: Predicting amino acid changes that affect protein function. Nucleic acids research. 2003;31(13):3812–4.

35. Lin W, Mandal S, Degen D, Liu Y, Ebright YW, Li S, et al. Structural basis of Mycobacterium tuberculosis transcription and transcription inhibition. Molecular cell. 2017;66(2):169–79.e8.

36. Morris GM, Huey R, Lindstrom W, Sanner MF, Belew RK, Goodsell DS, et al. AutoDock4 and AutoDockTools4: Automated docking with selective receptor flexibility. Journal of computational chemistry. 2009;30(16):2785–91.

37. Johansson MU, Zoete V, Michielin O, Guex N. Defining and searching for structural motifs using DeepView/Swiss-PdbViewer. BMC bioinformatics. 2012;13(1):173.

38. Fuhrmann J, Rurainski A, Lenhof HP, Neumann D. A new Lamarckian genetic algorithm for flexible ligand-receptor docking. Journal of computational chemistry. 2010;31(9):1911–8.

39. McEvoy CR, Cloete R, Müller B, Schürch AC, Van Helden PD, Gagneux S, et al. Comparative analysis of Mycobacterium tuberculosis pe and ppe genes reveals high sequence variation and an apparent absence of selective constraints. PloS one. 2012;7(4):e30593.

40. Joon M, Bhatia S, Pasricha R, Bose M, Brahmachari V. Functional analysis of an intergenic non-coding sequence within mce1 operon of M. tuberculosis. BMC microbiology. 2010;10(1):128.

41. Pang Y, Lu J, Wang Y, Song Y, Wang S, Zhao Y. Study of the rifampin monoresistance mechanism in Mycobacterium tuberculosis. Antimicrobial agents and chemotherapy. 2013;57(2):893–900.

42. Comas I, Borrell S, Roetzer A, Rose G, Malla B, Kato-Maeda M, et al. Whole-genome sequencing of rifampicin-resistant Mycobacterium tuberculosis strains identifies compensatory mutations in RNA polymerase genes. Nature genetics. 2012;44(1):106.

43. Li Q-j, Jiao W-w, Yin Q-q, Xu F, Li J-q, Sun L, et al. Compensatory mutations of rifampin resistance are associated with transmission of multidrug-resistant Mycobacterium tuberculosis Beijing genotype strains in China. Antimicrobial agents and chemotherapy. 2016;60(5):2807–12.

44. Telenti A, Philipp WJ, Sreevatsan S, Bernasconi C, Stockbauer KE, Wieles B, et al. The emb operon, a gene cluster of Mycobacterium tuberculosis involved in resistance to ethambutol. Nature medicine. 1997;3(5):567.

45. Sreevatsan S, Stockbauer KE, Pan X, Kreiswirth BN, Moghazeh SL, Jacobs WR, et al. Ethambutol resistance in Mycobacterium tuberculosis: critical role of embB mutations. Antimicrobial agents and chemotherapy. 1997;41(8):1677–81.

46. Shi D, Li L, Zhao Y, Jia Q, Li H, Coulter C, et al. Characteristics of embB mutations in multidrug-resistant Mycobacterium tuberculosis isolates in Henan, China. Journal of antimicrobial chemotherapy. 2011;66(10):2240–7.

47. Safi H, Sayers B, Hazbón MH, Alland D. Transfer of embB codon 306 mutations into clinical Mycobacterium tuberculosis strains alters susceptibility to ethambutol, isoniazid, and rifampin. Antimicrobial agents and chemotherapy. 2008;52(6):2027–34.

48. Cui Z, Li Y, Cheng S, Yang H, Lu J, Hu Z, et al. Mutations in the embC-embA region contribute to M. tuberculosis resistance to ethambutol. Antimicrobial agents and chemotherapy. 2014:AAC. 03285–14.

49. Sharma K, Gupta M, Pathak M, Gupta N, Koul A, Sarangi S, et al. Transcriptional control of the mycobacterial embCAB operon by PknH through a regulatory protein, EmbR, in vivo. Journal of bacteriology. 2006;188(8):2936–44.

50. Rastogi N, Labrousse V, Goh KS. In vitro activities of fourteen antimicrobial agents against drug susceptible and resistant clinical isolates of Mycobacterium tuberculosis and comparative intracellular activities against the virulent H37Rv strain in human macrophages. Current microbiology. 1996;33(3):167–75.

51. Shrivastava K, Garima K, Narang A, Bhattacharyya K, Vishnoi E, Singh RK, et al. Rv1458c: a new diagnostic marker for identification of Mycobacterium tuberculosis complex in a novel duplex PCR assay. Journal of medical microbiology. 2017;66(3):371–6.

52. Varma-Basil M, Garima K, Pathak R, Dwivedi SKD, Narang A, Bhatnagar A, et al. Development of a novel PCR restriction analysis of the hsp65 gene as a rapid method to screen for the Mycobacterium tuberculosis complex and nontuberculous mycobacteria in high-burden countries. Journal of clinical microbiology. 2013;51(4):1165–70.

53. Singh A, Kashyap VK. Specific and rapid detection of Mycobacterium tuberculosis complex in clinical samples by polymerase chain reaction. Interdisciplinary perspectives on infectious diseases. 2012;2012.

