## Supplementary figures and images for "Correlation of drug resistance with Single Nucleotide Variations through genome analysis and experimental validation in a multi-drug resistant clinical isolate of *M.tuberculosis*"

### Additional file 1

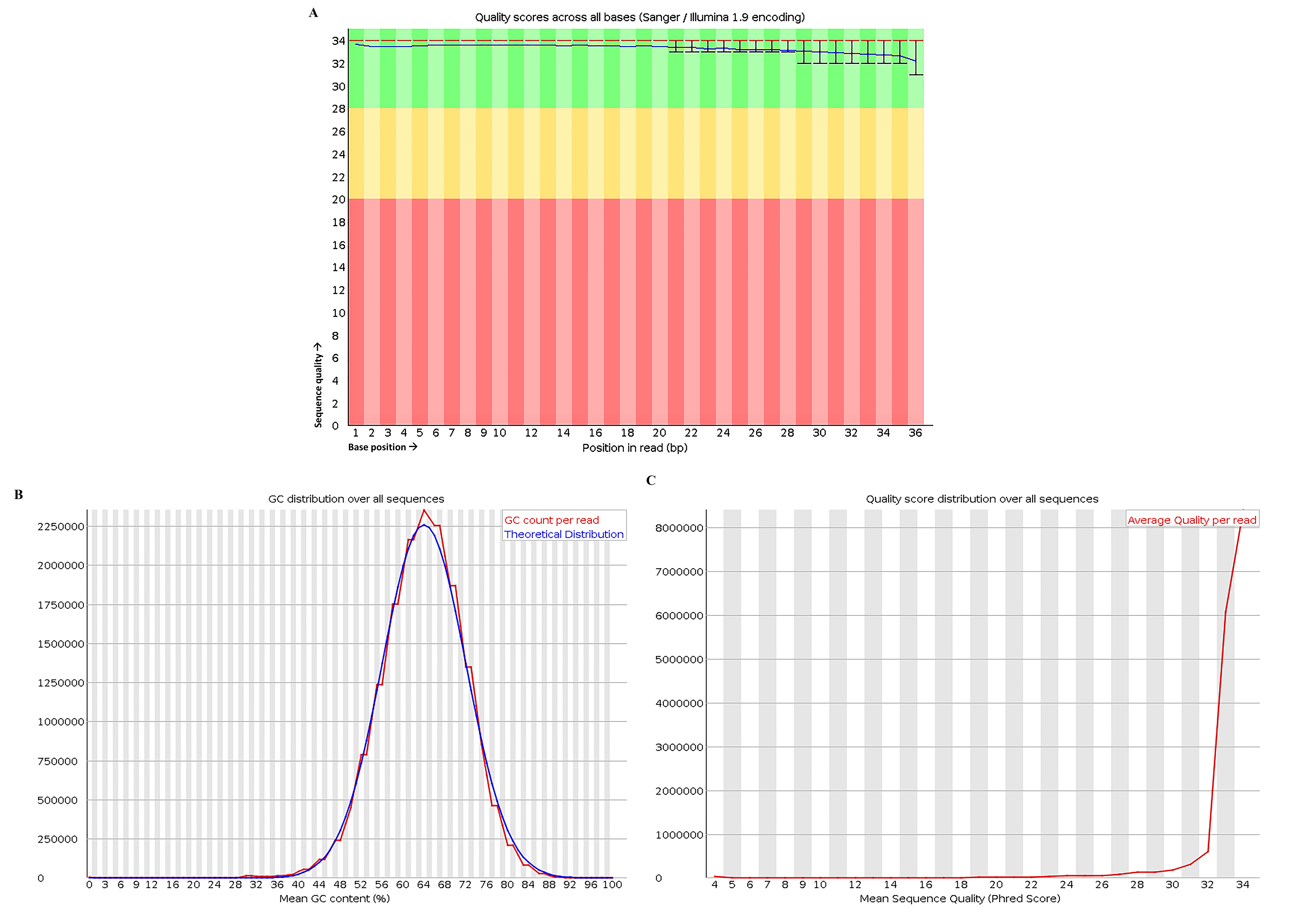

### Additional file 8

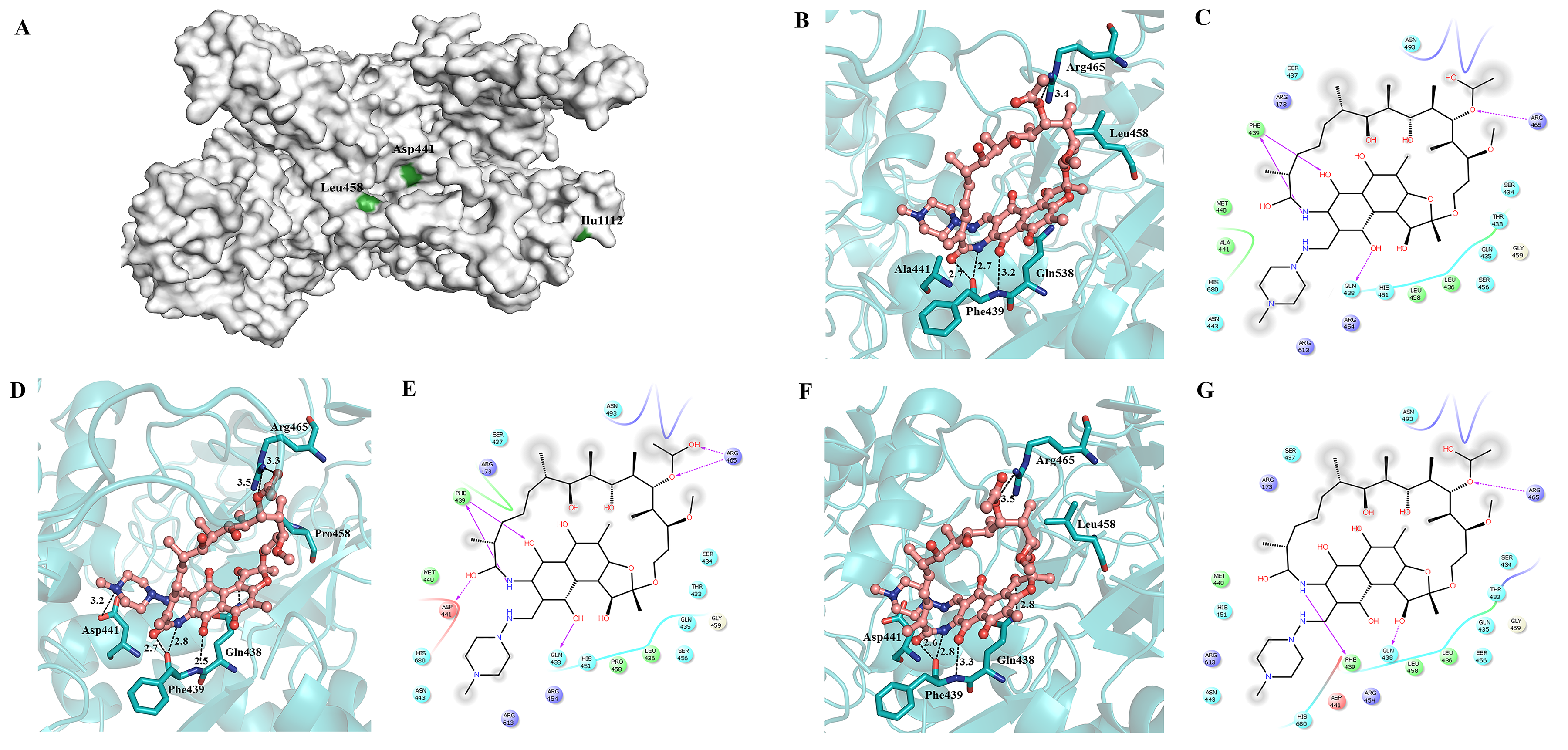
