## Additional file 3 for "Correlation of drug resistance with Single Nucleotide Variations through genome analysis and experimental validation in a multi-drug resistant clinical isolate of *M.tuberculosis*"

Intergenic Variations in VPCI591 (Type I)

| **Genomic Position of Variations** | **Wild Type** | **Variant** | **Genomic Position of Variations** | **Wild Type** | **Variant** |
| --- | --- | --- | --- | --- | --- |
| 1718 | C | A | 610120 | T | G |
| 1977 | A | G | 655986 | T | G |
| 5075 | C | T | 659341 | T | C |
| 7268 | C | T | 684290 | A | G |
| 13699 | T | C | 697808 | C | T |
| 26959 | C | G | 749968 | T | C |
| 27469 | A | G | 753668 | C | T |
| 34044 | T | C | 767414 | G | A |
| 46552 | C | G | 767609 | A | G |
| 47343 | G | A | 781395 | T | C |
| 66892 | C | G | 786137 | A | C |
| 68551 | A | G | 800357 | C | A |
| 79479 | T | C | 829719 | G | A |
| 80616 | C | G | 847995 | T | C |
| 97696 | T | C | 934230 | C | G |
| 133839 | C | T | 934611 | G | T |
| 139756 | T | C | 951142 | C | A |
| 139954 | A | C | 976043 | G | T |
| 196800 | G | C | 986463 | G | C |
| 226676 | G | A | 1067529 | C | T |
| 251635 | G | A | 1075279 | T | C |
| 251669 | T | C | 1077921 | C | T |
| 323306 | A | G | 1131464 | G | T |
| 333212 | G | C | 1148259 | A | G |
| 333292 | A | G | 1164336 | G | A |
| 409297 | A | G | 1174953 | A | C |
| 424250 | A | G | 1176926 | C | T |
| 439711 | T | C | 1200418 | A | G |
| 452288 | C | T | 1220570 | G | T |
| 459399 | A | C | 1224367 | T | C |
| 483935 | T | G | 1240894 | C | T |
| 503354 | G | C | 1262230 | C | T |
| 528354 | C | A | 1287081 | C | T |
| 534427 | T | A | 1287112 | T | C |
| 565655 | A | G | 1287160 | A | C |
| 1351172 | A | G | 2177073 | T | C |
| 1388644 | G | T | 2187363 | T | C |
| 1413148 | C | T | 2216571 | C | T |
| 1417554 | G | C | 2223293 | T | C |
| 1440771 | T | G | 2228967 | A | G |
| 1441334 | G | A | 2231486 | A | G |
| 1445977 | C | T | 2238930 | A | C |
| 1446005 | T | C | 2240062 | C | G |
| 1471659 | C | T | 2251999 | A | G |
| 1472362 | C | T | 2260094 | T | C |
| 1525160 | G | A | 2260100 | C | T |
| 1535643 | C | T | 2260525 | C | T |
| 1563686 | G | T | 2263278 | C | T |
| 1589234 | C | A | 2265059 | T | G |
| 1593347 | C | T | 2321358 | G | C |
| 1611283 | T | C | 2376425 | A | G |
| 1613035 | T | C | 2379743 | C | G |
| 1616831 | A | G | 2390299 | A | G |
| 1617833 | C | T | 2405619 | G | A |
| 1630279 | G | T | 2424864 | A | G |
| 1728615 | A | C | 2424925 | A | G |
| 1728837 | A | G | 2432170 | A | G |
| 1735903 | A | G | 2447539 | A | G |
| 1836286 | G | C | 2470485 | G | T |
| 1902378 | T | C | 2470591 | A | C |
| 1906336 | A | G | 2500892 | C | T |
| 1931470 | G | A | 2505919 | A | G |
| 1933988 | G | A | 2516567 | G | C |
| 1934234 | G | T | 2565061 | G | A |
| 1992715 | G | T | 2573756 | C | A |
| 2010614 | G | A | 2582324 | G | A |
| 2017560 | A | T | 2583803 | G | A |
| 2072409 | A | C | 2608488 | T | C |
| 2074754 | C | T | 2619271 | T | C |
| 2108890 | A | C | 2632500 | A | G |
| 2123169 | T | G | 2637088 | C | T |
| 2130529 | A | G | 2637541 | C | T |
| 2135870 | T | C | 2680658 | T | G |
| 2167489 | T | C | 2705603 | G | A |
| 2713795 | C | T | 3415332 | A | G |
| 2718852 | T | G | 3431407 | G | T |
| 2726051 | G | A | 3509091 | G | C |
| 2740693 | T | C | 3621423 | A | G |
| 2745310 | G | A | 3653386 | C | T |
| 2745739 | G | A | 3671843 | A | C |
| 2752698 | C | A | 3743549 | A | G |
| 2759534 | C | G | 3772616 | G | A |
| 2765565 | C | T | 3801551 | A | T |
| 2801147 | C | T | 3854899 | T | C |
| 2821709 | G | A | 3883467 | A | G |
| 2827984 | G | T | 3895727 | C | A |
| 2828019 | T | C | 3989107 | C | T |
| 2869242 | T | C | 4019103 | G | T |
| 2870386 | C | T | 4040719 | T | C |
| 2881244 | A | G | 4047653 | C | T |
| 2897871 | A | G | 4056416 | C | A |
| 2927939 | T | C | 4057081 | A | G |
| 2953307 | G | T | 4057367 | C | A |
| 2959324 | A | G | 4059904 | A | G |
| 2969006 | C | T | 4100975 | T | C |
| 3025431 | C | T | 4101018 | C | T |
| 3037377 | T | C | 4117161 | C | A |
| 3081599 | G | A | 4120983 | A | G |
| 3086788 | T | C | 4137190 | T | C |
| 3102196 | C | T | 4190532 | A | C |
| 3114814 | G | A | 4190596 | A | C |
| 3119513 | C | T | 4197189 | A | G |
| 3137237 | T | C | 4243229 | C | T |
| 3208522 | G | A | 4251077 | G | C |
| 3214329 | T | C | 4263279 | A | G |
| 3214481 | G | T | 4265593 | G | T |
| 3290535 | T | C | 4329782 | G | A |
| 3302683 | C | T | 4336597 | C | A |
| 3306594 | T | G | 4338732 | G | A |
| 3308606 | G | A | 4340234 | G | C |
| 3312942 | G | A | 4356690 | G | A |
| 3319485 | G | A | 4384007 | C | G |
| 3346121 | A | T | 4393178 | A | G |
| 3363338 | A | G | 4408920 | A | G |
| 3369869 | G | T | 4408923 | C | T |
| 3415051 | G | C | 4411016 | G | A |
