## Additional file 4 for "Correlation of drug resistance with Single Nucleotide Variations through genome analysis and experimental validation in a multi-drug resistant clinical isolate of *M.tuberculosis*"

**Additional file 4:** Intergenic Variations identified in VPCI591 (Type II)

| **Rv id** | **Function** | **Distance** | **SNV_pos** | **Distance** | **Function** | **Rv id** |
| --- | --- | --- | --- | --- | --- | --- |
| Rv0193c | Hypothetical | 105 | 226676 | 201 | Multidrug efflux pump | Rv0194 |
| Rv0439c | Probable dehydrogenase | 40 | 528354 | 253 | Chaperonin 2 GroEL2 | Rv0440 |
| Rv0653c | Transcriptional regulator TetR | 39 | 749968 | 31 | Probable dioxygenase | Rv0654 |
| Rv0837c | Hypothetical | 199 | 934230 | 489 | Lipoprotein LpqR | Rv0838 |
| Rv0853c | Pyruvate decarboxylase | 24 | 951142 | 40 | Hypothetical | Rv0854 |
| Rv0876c | Hypothetical | 106 | 976043 | 31 | Hypothetical | Rv0877 |
| Rv0966c | Hypothetical | 86 | 1077921 | 53 | Cu sensitive operon repressor | Rv0967 |
| Rv1075c | Hypothetical | 48 | 1200418 | 348 | Lipase LipU | Rv1076 |
| Rv1435c | Hypothetical | 85 | 1613035 | 271 | GAPDH | Rv1436 |
| Rv1706c | Hypothetical | 110 | 1933988 | 493 | PPE23 | Rv1706A |
| Rv1759c | PE-PGRS | 138 | 1992715 | 437 | Triacylglycerol synthase | Rv1760 |
| Rv1776c | Transcriptional regulator | 59 | 2010614 | 41 | Cytochrome P450 | Rv1777 |
| Rv1872c | Lactate dehydrogenase | 18 | 2123169 | 4 | Hypothetical | Rv1873 |
| Rv1886c | Secreted antigen 85-B | 3 | 2135870 | 387 | Hypothetical | Rv1887 |
| Rv1924c | Hypothetical | 143 | 2177073 | 13 | Acyl-CoA ligase | Rv1925 |
| Rv2005c | Hypothetical | 116 | 2251999 | 2 | Trehalose-phosphatase | Rv2006 |
| Rv2115c | Proteasome ATPase | 135 | 2376425 | 145 | Lipoprotein LppK | Rv2116 |
| Rv2205c | Hypothetical | 22 | 2470485 | 136 | Hypothetical | Rv2206 |
| Rv2293c | Hypothetical | 29 | 2565061 | 265 | Aminotransferase | Rv2294 |
| Rv2333c | Drug efflux protein | 167 | 2608488 | 307 | Cysteine synthase | Rv2334 |
| Rv2421c | Nucleotide adenylyltransferase | 44 | 2718852 | 230 | Hypothetical | Rv2422 |
| Rv2457c | ATP-binding subunit ClpX | 46 | 2759534 | 244 | Cysteine methyltransferase | Rv2458 |
| Rv2505c | Hypothetical | 113 | 2821709 | 2 | Transcriptional regulator | Rv2506 |
| Rv2572c | Aspartate--tRNA ligase | 68 | 2897871 | 84 | Hypothetical | Rv2573 |
| Rv2712c | Hypothetical | 103 | 3025431 | 9 | Pyridine transhydrogenase | Rv2713 |
| Rv2779c | Transcriptional regulator | 34 | 3086788 | 31 | Alanine dehydrogenase | Rv2780 |
| Rv2830c | Antitoxin VapB22 | 13 | 3137237 | 33 | Enoyl-CoA hydratase | Rv2831 |
| Rv2904c | 50S ribosomal protein | 76 | 3214329 | 298 | Alanine rich lipoprotein | Rv2905 |
| Rv2955c | Hypothetical | 61 | 3308606 | 61 | Hypothetical | Rv2956 |
| Rv2988c | Iisopropylmalate dehydratase | 46 | 3346121 | 25 | Transcriptional regulator | Rv2989 |
| Rv3271c | Membrane protein | 39 | 3653386 | 61 | Hypothetical | Rv3272 |
| Rv3694c | Hypothetical | 76 | 4137190 | 15 | Membrane protein | Rv3695 |
