## Additional file 5 for "Correlation of drug resistance with Single Nucleotide Variations through genome analysis and experimental validation in a multi-drug resistant clinical isolate of *M.tuberculosis*"

**Additional file 5:** Invariant genes in global clinical isolates (including VPCI591)

| **Gene Id** | **Known Function** |
| --- | --- |
| Rv0334 | Alpha-D-glucose-1-phosphate thymidylyltransferase RmlA (dTDP-glucose synthase) |
| Rv0491 | Two component sensory transduction protein RegX3 |
| Rv0550c | Possible antitoxin VapB3 |
| Rv0581 | Possible antitoxin VapB26 |
| Rv0603 | Possible exported protein |
| Rv0661c | Possible toxin VapC7 |
| Rv0664 | Possible antitoxin VapB8 |
| Rv0666 | Possible membrane protein |
| Rv0700 | 30S ribosomal protein S10 RpsJ |
| Rv0718 | 30S ribosomal protein S8 RpsH |
| Rv0720 | 50S ribosomal protein L18 RplR |
| Rv0793 | Possible monooxygenase |
| Rv0795 | Putative transposase for insertion sequence element IS6110 |
| Rv0796 | Putative transposase for insertion sequence element IS6110 |
| Rv0865 | Probable molybdopterin biosynthesis Mog protein |
| Rv0871 | Probable cold shock-like protein B CspB |
| Rv0900 | Possible membrane protein |
| Rv0947 | Probable mycolyl transferase |
| Rv0961 | Probable integral membrane protein |
| Rv1024 | Possible conserved membrane protein |
| Rv1042c | Putative transposase for insertion element |
| Rv1055 | Possible integrase |
| Rv1103c | Possible antitoxin MazE3 |
| Rv1149 | Possible transposase |
| Rv1150 | Possible transposase |
| Rv1195 | PE family protein PE13 |
| Rv1220c | Probable methyltransferase |
| Rv1237 | Probable sugar-transport integral membrane protein ABC transporter SugB |
| Rv1369c | Probable transposase |
| Rv1370c | Putative transposase for insertion sequence element IS6111 |
| Rv1379 | Probable pyrimidine operon regulatory protein PyrR |
| Rv1398c | Possible antitoxin VapB10 |
| Rv1440 | Probable protein-export membrane protein (translocase subunit) SecG |
| Rv1473 | Probable macrolide-transport ATP-binding protein ABC transporter |
| Rv1580c | Probable PhiRv1 phage protein |
| Rv1581c | Probable PhiRv1 phage protein |
| Rv1582c | Probable PhiRv1 phage protein |
| Rv1583c | Probable PhiRv1 phage protein |
| Rv1584c | Probable PhiRv1 phage protein |
| Rv1586c | Probable PhiRv1 integrase |
| Rv1721c | Possible antitoxin VapB12 |
| Rv1756c | Putative transposase for insertion sequence element IS6110 |
| Rv1757c | Putative transposase for insertion sequence element IS6110 |
| Rv1763 | Putative transposase for insertion sequence element IS6110 |
| Rv1764 | Putative transposase for insertion sequence element IS6110 |
| Rv1792 | ESAT-6 like protein EsxM |
| Rv1799 | Probable lipoprotein LppT |
| Rv1853 | Probable urease accessory protein UreD |
| Rv1994c | Metal sensor transcriptional regulator CmtR |
| Rv2034 | ArsR repressor protein |
| Rv2055c | 30S ribosomal protein S18 RpsR2 |
| Rv2060 | Possible conserved integral membrane protein |
| Rv2094c | Sec-independent protein translocase membrane-bound protein TatA |
| Rv2098c | PE-PGRS family protein PE_PGRS36 |
| Rv2099c | PE family protein PE21 |
| Rv2105 | Putative transposase for insertion sequence element IS6110 |
| Rv2106 | Probable transposase |
| Rv2149c | Conserved protein YfiH |
| Rv2167c | Probable transposase |
| Rv2168c | Putative transposase for insertion sequence element IS6110 |
| Rv2238c | Probable peroxiredoxin AhpE |
| Rv2278 | Putative transposase for insertion sequence element IS6110 |
| Rv2279 | Putative transposase for insertion sequence element IS6110 |
| Rv2354 | Putative transposase for insertion sequence element IS6110 |
| Rv2355 | Putative transposase for insertion sequence element IS6110 |
| Rv2403c | Probable conserved lipoprotein LppR |
| Rv2431c | PE family protein PE25 |
| Rv2441c | 50S ribosomal protein L27 RpmA |
| Rv2445c | Probable nucleoside diphosphate kinase NdkA |
| Rv2479c | Probable transposase- Involved in the transposition of the insertion sequence IS6110 |
| Rv2480c | Possible transposase for insertion sequence element IS6110 (fragment) |
| Rv2520c | Possible conserved membrane protein |
| Rv2526 | Possible antitoxin VapB17 |
| Rv2539c | Shikimate kinase AroK |
| Rv2550c | Possible antitoxin VapB20 |
| Rv2581c | Possible glyoxalase II |
| Rv2639c | Probable conserved integral membrane protein |
| Rv2642 | Possible transcriptional regulatory protein (probably ArsR-family) |
| Rv2648 | Probable transposase for insertion sequence element IS6110 (fragment) |
| Rv2649 | Probable transposase for insertion sequence element IS6110 |
| Rv2654c | Possible PhiRv2 prophage protein |
| Rv2656c | Possible PhiRv2 prophage protein |
| Rv2657c | Probable PhiRv2 prophage protein |
| Rv2658c | Possible prophage protein |
| Rv2760c | Possible antitoxin VapB42 |
| Rv2806 | Possible membrane protein |
| Rv2814c | Probable transposase- Involved in the transposition of the insertion sequence IS6110 |
| Rv2815c | Probable transposase- Involved in the transposition of the insertion sequence IS6110 |
| Rv2829c | Possible toxin VapC22 |
| Rv2872 | Possible toxin VapC43 |
| Rv2906c | Probable tRNA (guanine-N1)-methyltransferase TrmD |
| Rv2965c | Probable phosphopantetheine adenylyltransferase KdtB |
| Rv3002c | Probable acetolactate synthase (small subunit) IlvN (acetohydroxy-acid synthase) |
| Rv3013 | Conserved protein |
| Rv3058c | Possible transcriptional regulatory protein (probably TetR-family) |
| Rv3147 | Probable NADH dehydrogenase I (chain C) NuoC (NADH-ubiquinone oxidoreductase chain C) |
| Rv3149 | Probable NADH dehydrogenase I (chain E) NuoE (NADH-ubiquinone oxidoreductase chain E) |
| Rv3155 | Probable NADH dehydrogenase I (chain K) NuoK (NADH-ubiquinone oxidoreductase chain K) |
| Rv3184 | Probable transposase for insertion sequence element IS6110 (fragment) |
| Rv3185 | Probable transposase- Involved in the transposition of the insertion sequence IS6110 |
| Rv3186 | Probable transposase for insertion sequence element IS6110 (fragment) |
| Rv3187 | Probable transposase- Involved in the transposition of the insertion sequence IS6110 |
| Rv3216 | GCN5-related N-acetyltransferase, pseudogene |
| Rv3287c | Anti-sigma factor RsbW |
| Rv3321c | Possible antitoxin VapB44 |
| Rv3325 | Probable transposase for insertion sequence element IS6110 (fragment) |
| Rv3326 | Probable transposase- Involved in the transposition of the insertion sequence IS6110 |
| Rv3352c | Possible oxidoreductase |
| Rv3380c | Probable transposase- Involved in the transposition of the insertion sequence IS6110 |
| Rv3381c | Probable transposase for insertion sequence element IS6110 (fragment) |
| Rv3414c | Probable alternative RNA polymerase sigma-D factor SigD |
| Rv3453 | Possible conserved transmembrane protein |
| Rv3461c | 50S ribosomal protein L36 RpmJ |
| Rv3474 | Possible transposase for insertion element IS6110 (fragment) |
| Rv3475 | Possible transposase for insertion element IS6110 [second part] |
| Rv3503c | Probable ferredoxin FdxD |
| Rv3574 | Transcriptional regulatory protein KstR |
| Rv3615c | ESX-1 secretion-associated protein EspC |
| Rv3622c | PE family protein PE32 |
| Rv3844 | Possible transposase |
| Rv3851 | Possible membrane protein |
| Rv3875 | 6 kDa early secretory antigenic target EsxA (ESAT-6) |
| Rv3924c | 50S ribosomal protein L34 RpmH |
