## Additional file 6 for "Correlation of drug resistance with Single Nucleotide Variations through genome analysis and experimental validation in a multi-drug resistant clinical isolate of *M.tuberculosis*"

**Additional file 6:** Unique genic variations in VPCI591

| **Gene Name** | **Rv id** | **Nucleotide Change** | **Type of Change** | **Amino Acid Change** | **SNV position in genome** |
| --- | --- | --- | --- | --- | --- |
| whiB5 | Rv0022c | T293C | nonsynonymous | I98T | 27150 |
| nrp | Rv0101 | A3421G | nonsynonymous | I1141V | 113421 |
| PE3 | Rv0159c | C176G | nonsynonymous | A59G | 188664 |
| PPE8 | Rv0355c | C8178A | nonsynonymous | N2726K | 426502 |
| pks6 | Rv0405 | T3755G | nonsynonymous | L1252R | 489485 |
| ctpH | Rv0425c | G52T | nonsynonymous | G18C | 515270 |
| sigK | Rv0445c | C24G | nonsynonymous | S8R | 534373 |
| proC | Rv0500 | G316A | nonsynonymous | G106S | 590398 |
| rpoB | Rv0667 | A1304C | nonsynonymous | D435A | 761110 |
| rpoB | Rv0667 | C3318G | nonsynonymous | I1106M | 763124 |
| xylB | Rv0729 | C326T | nonsynonymous | S109L | 821832 |
| purM | Rv0809 | C23A | nonsynonymous | P8H | 903747 |
| PE7 | Rv0916c | C41T | nonsynonymous | A14V | 1021603 |
| uvrD1 | Rv0949 | C2000T | nonsynonymous | P667L | 1060259 |
| lgt | Rv1614 | T923C | nonsynonymous | V308A | 1814093 |
| uvrA | Rv1638 | C244T | nonsynonymous | P82S | 1843984 |
| pks8 | Rv1662 | T3419G | nonsynonymous | V1140G | 1885122 |
| yrbE3B | Rv1965 | G160A | nonsynonymous | A54T | 2208666 |
| cobN | Rv2062c | G2989T | nonsynonymous | A997S | 2317765 |
| pyrD | Rv2139 | G472A | nonsynonymous | D158N | 2399191 |
| pyrD | Rv2139 | C1063T | stopgain | Q355X | 2399782 |
| qcrC | Rv2194 | T152C | nonsynonymous | V51A | 2457704 |
| accD6 | Rv2247 | T712C | nonsynonymous | S238P | 2521454 |
| cyp121 | Rv2276 | G949A | nonsynonymous | E317K | 2548697 |
| aroB | Rv2538c | C587A | nonsynonymous | P196Q | 2862090 |
| speE | Rv2601 | C967T | stopgain | Q323X | 2929354 |
| PPE42 | Rv2608 | C1452A | stopgain | Y484X | 2936497 |
| smc | Rv2922c | G641A | nonsynonymous | R214H | 3237166 |
| ppsB | Rv2932 | G412A | nonsynonymous | G138S | 3251483 |
| mas | Rv2940c | G5525A | nonsynonymous | G1842E | 3277191 |
| recG | Rv2973c | C653T | nonsynonymous | P218L | 3329294 |
| gltX | Rv2992c | A623C | nonsynonymous | K208T | 3349655 |
| pgmA | Rv3068c | C682T | nonsynonymous | R228W | 3432941 |
| lat | Rv3290c | G161A | nonsynonymous | R54Q | 3671634 |
| lpqD | Rv3390 | C412T | nonsynonymous | P138S | 3805276 |
| phyA | Rv3397c | G262T | nonsynonymous | A88S | 3814737 |
| PPE57 | Rv3425 | G328A | nonsynonymous | A110T | 3842566 |
| PPE58 | Rv3426 | C470A | nonsynonymous | P157Q | 3843505 |
| panD | Rv3601c | C176T | nonsynonymous | A59V | 4044106 |
| dppA | Rv3666c | C878T | nonsynonymous | A293V | 4106207 |
| embB | Rv3795 | A1512C | nonsynonymous | E504D | 4248025 |
| PPE68 | Rv3873 | A229G | nonsynonymous | T77A | 4351303 |
