## Additional file 7 for "Correlation of drug resistance with Single Nucleotide Variations through genome analysis and experimental validation in a multi-drug resistant clinical isolate of *M.tuberculosis*"

**Additional file 6:** Inter-genic variations unique to VPCI591

| **Adjacent gene** | **Genomic SNV position** | **Adjacent gene** | **Wild type** | **Variant**  **type** |
| --- | --- | --- | --- | --- |
| fadD5 | 196800* | yrbE1A | G | C |
| Rv0966c | 1077921 | csoR | C | T |
| Rv0102 | 1131464 | Pks16 | G | T |
| Rv0151c | 1174953 | Rv0152 | A | C |
| Rv1115 | 1240894 | Rv1116 | C | T |
| Rv1245c | 1388644 | relE | G | T |
| PPE23 | 1934234 | Rv1706A | G | T |
| Wag32 | 1992715 | Rv1760 | G | T |
| Wag31 | 2405619 | Rv2146c | G | A |
| Rv2644c | 2969006 | valT | C | T |
| Rv2792c | 3102196 | truB | C | T |
| rplS | 3214329 | lppw | T | C |
| Rv3387 | 3801551** | PE_PGRS52 | A | T |
| embC | 4243229*** | embA | C | T |
| Rv3796 | 4251077 | fadE35 | G | C |
| Rv3863 | 4340234 | espE | G | C |

* SNVs mapping in the regulatory region in Mce1 operon (40), **SNV within IS1560 element, ***SNV mapping in the mutation hotspot and the adjacent regions are known for drug resistance to ethambutol (41).
